# Single-Molecule Kinetic Exploration of Functional Substates in an Evolving Phosphotriesterase

**DOI:** 10.1101/2025.04.14.648755

**Authors:** Morito Sakuma, Dhani Ram Mahato, Ferran Feixas, Colin J. Jackson, Eiji Nakata, Sílvia Osuna, Nobuhiko Tokuriki

## Abstract

Enzymes achieve catalysis by dynamically sampling diverse conformational states. Beyond this plasticity, individual enzyme molecules occupy metastable substates, forming an ensemble of functional substates within a population. Since shifts in functional substate dynamics drive phenotypic variation, their evolutionary trajectories are central to the emergence of new functions. However, the challenge of measuring functional substates has hindered our understanding of their role in enzyme evolution and the optimization of conformational substates. Here, we address this gap by investigating how functional and conformational substates were modulated during enzyme functional transitions, using single-molecule kinetic assays and molecular dynamics simulations. We analyzed wild-type phosphotriesterase (PTE) and 18 evolved variants that transitioned from the native PTE to promiscuous arylesterase (AE) activity. Our findings reveal that evolutionary transitions reshape functional and conformational substate landscapes: PTE-specialized variants exhibit broader substate distributions, whereas AE-specialized variants display more uniform substates. These results provide the first direct evidence that enzyme evolution is accompanied by coordinated shifts in functional and conformational substate equilibria, optimizing both for the enzyme’s catalytic efficiency. This work highlights the power of single-molecule techniques in uncovering how heterogeneous enzyme populations navigate substate transitions and, ultimately, how these transitions shape enzyme evolvability.

## Introduction

The inherent flexibility of protein structures in solution facilitates sampling a broad spectrum of conformational substates [1, 2]. Such distinct structural arrangements and dynamic transitions among substates are central to models of enzyme-substrate interactions in the catalytic cycle, such as induced fit and conformational selection [3–9]. Beyond their dynamic nature, enzymes can exist as metastable conformational substates, giving rise to functional substates, *i.e*., individual enzyme molecules in different conformational substates exhibit distinct catalytic efficiencies and specificities despite having an identical sequence. Various factors can contribute to the emergence of functional substates, including alternative folding, distinct ligand or cofactor binding, and oligomerization [10–16]. Given that an ensemble of functional substates and its shifting equilibrium in enzyme population directly link to phenotypic variation [17], understanding and controlling these states are critical challenges in enzyme engineering and design.

Current evolutionary theory suggests that mutations can shift the population distribution of the substates, favoring those that enhance promiscuous or specific activity [18–21]. This underscores the importance of developing advanced approaches to gain a deeper understanding of the heterogeneous nature of enzyme populations. Structural analysis, including NMR, has demonstrated a transition between active and inactive conformational states in enzyme populations [22, 23]. Advanced kinetic methods, particularly single-molecule kinetic assays, have provided direct insights into enzyme behavior at the molecular level, revealing previously cryptic aspects of conformational and functional states [24–26]. By tracking hundreds to thousands of individual enzymatic reactions, several studies have uncovered the presence of distinct functional substates that persist over minutes, suggesting long-lived functional heterogeneity in enzyme populations [27–32]. Intriguingly, although only a handful of enzymes have been tested, variants that sample more diverse functional substates display greater catalytic promiscuity, exhibiting activity across a broader range of potential substrates [33, 34]. These observations suggest that enzyme ensemble includes molecules with distinct reactivities toward new substrates. Thus, further investigation into functional substates, especially how they are altered during enzyme evolution and their relationship to multifunctionality and conformational substates, could provide deeper insights into enzyme evolvability.

Here, we applied single-molecule kinetic assays along with protein conformational dynamics characterization across a series of enzyme variants that represent a complete transition of enzyme functions (**Fig. 1**). We aim to uncover the link between the distributions of functional and conformational substates (**Fig. 1**). As a model system, we employed bacterial phosphotriesterase (PTE), a Zn^2+-^dependent metalloenzyme that is one of the best-studied for its evolutionary dynamics and is among the most active enzymes reported [35–38]. Previously, we evolved the wild-type PTE (R0) to enhance its promiscuous arylesterase (AE) activity over 22 rounds of directed evolution (forward evolution) and generated a highly efficient and specialized AE enzyme (R22) [35]. Subsequently, 12 rounds of evolution from R22 were performed to restore the PTE activity (reverse evolution), yielding neoPTE (or revR12), a bifunctional enzyme exhibiting high efficiency for both PTE and AE activities [36]. These overall trajectories generated 37 variants with more than a 10^8^-fold change in specificity between PTE and AE activities (**Fig. S1**) [35, 36, 38]. Structural characterizations unveiled the molecular mechanisms underlying the functional transitions [37], providing an excellent system to explore the existence and role of functional substates during enzyme evolution. We measured the single-molecule kinetics of wild-type and 18 PTE variants across the trajectories for both PTE and AE activities. In this study, we characterized changes in their functional substates for both original and evolved substrates. Through this analysis, we establish that the PTE optimized functional substates for each function during enzyme evolution, which were coupled with the optimization of local structures essential for each function. Thus, this work uncovers the cryptic role of static enzyme heterogeneity in enzyme evolution.

**Figure 1.**
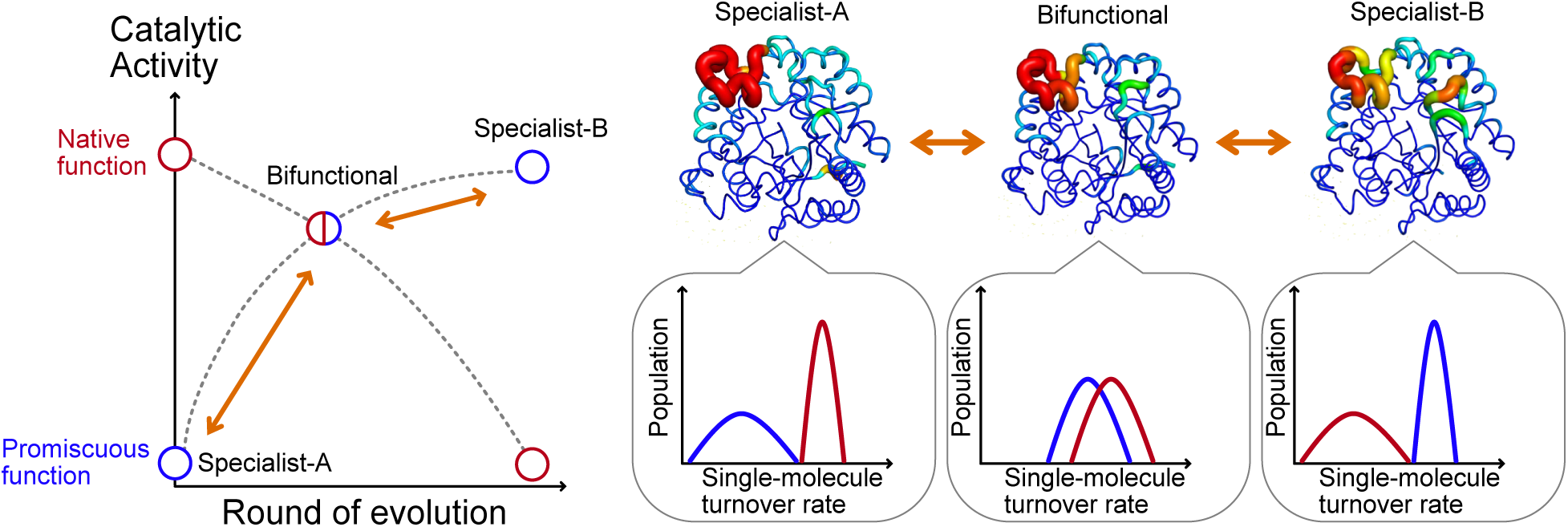
Protein evolution at the single-molecule level. Enzymes can evolve such that their promiscuous functions increase to the same level as their original native function (neofunctionalization). Through this evolutionary process, a bifunctional intermediate may arise, exhibiting comparable and significant activity in both its native and promiscuous functions. One hypothesis for the molecular basis of this neofunctionalization is that changes in conformational dynamics during the evolutionary trajectory generate heterogeneous functional substates that could catalyze reactions with both the native and new substrate or develop different substates depending on functions. To test this, we have measured the distribution of conformational and functional substates across an evolutionary trajectory to clarify their correlation.

## Results & Discussion

### Development of a single-molecule kinetic assay of PTE

We opted for a single-molecule kinetic assay of PTE variants using resorufin-diethylphosphate (RDP) and resorufin-butyrate (RB) for measuring functional substates (**Fig. 2A**). These substrates have relatively similar leaving group (resorufin), comparable to that of the original substrates used in the directed evolution (naphthol; **Fig. 2A** and **Fig. S1A**) [35, 36, 38]. We confirmed that RDP and RB largely recapitulated the changes in the activity observed during the directed evolution in an ensemble kinetic assay (**Fig. 2B**, **Fig. S1B,** and **Table S1**). R0 displayed high PTE activity (*k*_cat_/*K*_M_ = 1.3 × 10^8^ M^-1^ s^-1^) and low AE activity (*k*_cat_/*K*_M_ = 3.6 × 10^4^ M^-1^ s^-1^). Throughout evolution, these activities gradually reversed, with R6 displaying bifunctionality and high activities toward both substrates (*k*_cat_/*K*_M_ for RDP and RB were 3.5 × 10^6^ and 2.5 × 10^7^ M^-1^ s^-1^, respectively). Eventually, the last variant in the forward evolution, R22, became highly specific to AE activity (*k*_cat_/*K*_M_ = 3.5 × 10^7^ M^-1^ s^-1^ for RB *vs.* 1.4 × 10^3^ M^-1^ s^-1^ for RDP). In the reverse evolution, PTE activity was restored while AE activity was largely unaffected; consequently, revR12 exhibited bifunctional features (*k*_cat_/*K*_M_ for RDP and RB were 1.2 × 10^7^ and 1.1 × 10^7^ M^-1^ s^-1^, respectively).

**Figure 2.**
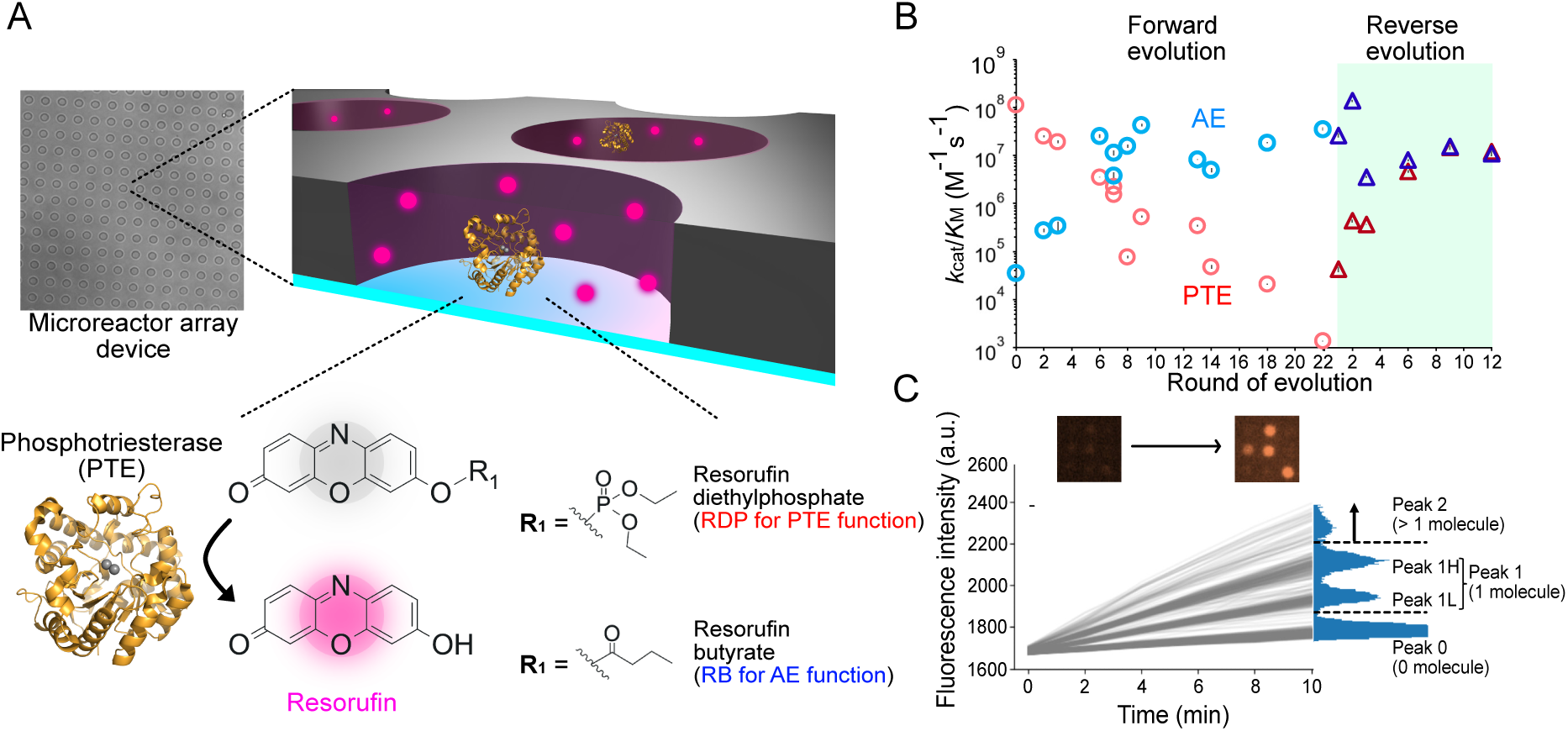
Single-molecule kinetic assay with a microreactor array device. (A) Single PTE molecules were trapped in the microreactor array. PTE hydrolyzed resorufin-based substrates (RDP and RB) and generated the fluorescent product (resorufin). (B) Bulk ensemble kinetic assay of PTE and AE activities. *k*_cat_/*K*_M_ (M^-1^ s^-1^) of purified wild-type PTE-R0 and its variants were measured with RDP (Red circles and triangles) and RB (blue circles and triangles) substrates (mean ± *SE*, n = 3). Forward and reverse evolution trajectories are shown as circles and triangles, respectively. (C) Time-course measurement of a typical catalytic reaction in the microreactor array. Microscope image was taken at 2-minute intervals for 10 minutes (example in top panel). The histogram of the fluorescence intensity distribution typically presents three peaks: peak 0 (enzyme-free reactors), peak 1 (single enzyme activity), and peak 2 (two or more enzyme molecules per reactor). For peak 1, the majority of enzyme variants showed a bimodal distribution with distinct low and high activity peaks (peak 1L and peak 1H). The dotted lines indicate thresholds to exclude peak 0 and peak 2 from the analysis of single-molecule activity.

We established single-molecule kinetic assays for RDP and RB hydrolysis using a microreactor array device (**Fig. 2A** and see **Supplementary Methods**) [32, 34, 39]. Each device displays over 30,000 reactors, into which an assay solution with highly diluted enzyme and 200 μM substrate was applied. This allowed the reaction rates of more than 700 single-enzyme reactions to be monitored without leakage of enzyme and substrates (**Fig. S2A** and **Fig. S3**). The histograms of fluorescence intensity at 10 min typically showed three peaks (peak 0, 1, and 2) (**Fig. 2C**). Peak 0 corresponds to reactors without enzyme molecules, peak 1 to single-molecule activity, and peak 2 represents reactors with two or more enzyme molecules. The relative populations of these peaks were consistent with a Poisson distribution [32, 34, 39], with the majority of the reactors falling into peak 0, followed by peak 1 and peak 2 (79%, 18%, and 2% of molecules belonged to peak 0, peak 1, and peak 2 when 11 was 0.23) (see **Supplementary Methods**).

We measured single enzyme kinetics of wild-type and 18 PTE variants across the laboratory evolution trajectories (**Fig. S4**). 14 of them were measured with both RDP and RB substrates while three variants (R0, R2, R3) were measured with only RDP, and two (R18, R22) were measured with only RB due to the detection limit of the single-molecule kinetic assay. We followed the reaction over time and found that each reactor with a single enzyme molecule showed a linear increase in fluorescence intensity for at least 10 minutes, which is consistent with previous studies with other enzyme systems [30–34, 39]. This suggests that each enzyme molecule retains a metastable functional substate (see **Supplementary Methods**). We calculated catalytic activity as the mean turnover rate using a calibration curve that correlates the concentration of resorufin dye with its fluorescence intensity in the microreactor array (**Fig. S5B**). The average catalytic activities observed in the single-molecule kinetic assay were consistent with those in the ensemble kinetic assay, indicating that the environment of the device and the method of measurement did not affect the function of PTEs or introduce any experimental artifacts (**Fig. S6A**).

### Single-molecule kinetic assay revealed a bimodal activity distribution

The time- course measurements of the various reactors identified peak 0 and peak 2 populations, where either zero or more than two enzyme molecules were present, respectively. However, it also revealed that peak 1 exhibited a bimodal distribution that could be separated into two distinct populations (**Fig. 2C** and **Fig. S4**), and this distribution was not associated with peak 2 according to Poisson statistics (see **Supplementary Methods**); we refer to them as peak 1H and peak 1L. Notably, the bimodal distribution changes for each variant along the trajectory (**Fig. S7A**). Thus, rather than a functionally relevant difference, it appears that this “splitting” of the peak 1 activity into two populations may be a systematic artifact of the experiment.

To investigate further, we first hypothesized that this could represent an equilibrium between two stable conformational substates with different activities (**Fig. 3A**). However, subsequent analysis revealed data that contradicted this hypothesis. The turnover rate ratios between peak 1L and peak 1H were largely consistent for most variants, where the turnover rate of peak 1H was approximately 1.6 times that of peak 1L (**Fig. S7B**). In addition, the population ratios between peak 1H and peak 1L for both PTE and AE functions followed a similar trend within the same variant (**Fig. S7C**). Finally, we directly assessed possible interconversion between the two peaks by employing reaction solution exchange experiments (**Fig. S2B** and see **Supplementary Methods**). This technique allows us to exchange the reaction solutions in the microreactors and measure the activities of identical enzyme molecules multiple times without impairing their activity (**Fig. S8B-C**) [34, 39]. We measured the same activity twice ((1^st^, 2^nd^) = (PTE, PTE) or (AE, AE)) and two different reactions ((1^st^, 2^nd^) = (PTE, AE)) and analyzed the transition between two peaks (**Fig. 3B-C**, **Fig. S8A**, and **Fig. S9**). We identified two dense regions in the density scatter plots of the first and second reactions (cluster 1 and cluster 2 in **Fig. 3B-C**, **Fig. S8A**, and **Fig. S9**). Most enzyme molecules were found within these regions, regardless of the reaction: the population ratios of cluster 1 and cluster 2 in the (PTE, PTE), (AE, AE), and (PTE, AE) measurements were 84 ± 4%, 84 ± 4%, and 81 ± 6% across all measurements, respectively (**Table S2**). Moreover, when we measured the same function twice, both clusters (1 and 2) exhibited a diagonally elongated distribution in the scattered plots. Spearman’s correlation coefficients (ρ) between two reactions, (PTE, PTE) and (AE, AE), in R6 were (cluster 1, cluster 2) = (0.40 ± 0.01, 0.35 ± 0.02, n = 2) and (cluster 1, cluster 2) = (0.33 ± 0.08, 0.38 ± 0.05) (mean ± *SE*, n = 2), respectively (**Fig. 3B-C**). Similar ρ values were also observed in various PTE variants (**Table S2**): (PTE, PTE) and (AE, AE) in all measured variants were (cluster 1, cluster 2) = (0.30 ± 0.02, 0.33 ± 0.03) and (0.34 ± 0.02, 0.36 ± 0.03) (mean ± *SE*), which are lower than the upper limit of the assay system (ρ_upper limit_ = 0.42 ± 0.05 (mean ± *SE*, n = 2) (see **Supplementary Methods**)). In addition, positive ρ values were also observed even though two different reactions (PTE, AE) were measured ((cluster 1, cluster 2) = (0.23 ± 0.02, 0.22 ± 0.02) (**Table S2)**. This suggests that enzyme molecules sample neighboring functional substates independent of their specific function in this experimental time scale. Altogether, these results suggest that there is no interconversion between the two peaks and that these are not different conformational substates but are static populations that stably exist.

**Figure 3.**
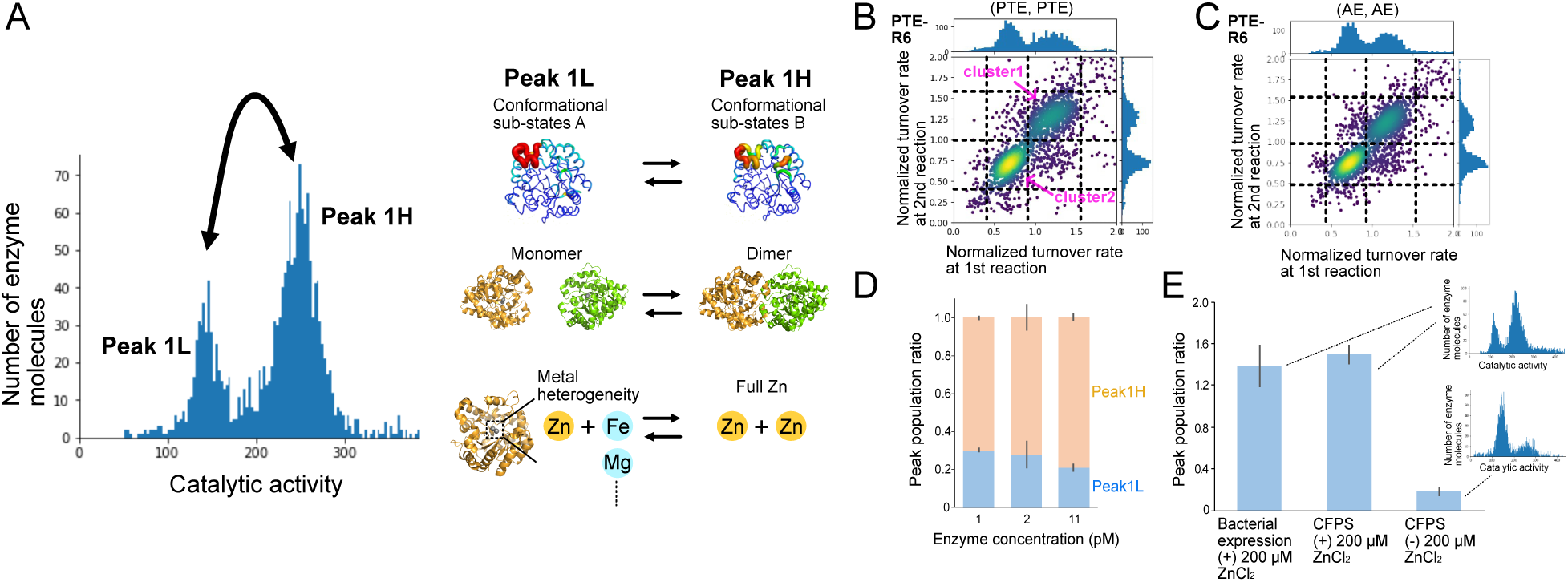
Molecular causes of the bimodal distribution in the single-molecule kinetic assay. (A) Three possible causes of the bimodal distribution (peak 1L and peak 1H): two different conformational substates, monomer-dimer transition, and metal heterogeneity in the active site. (B), (C) Reaction solution exchange experiment to measure the transition between peaks. PTE- R6 activity was measured with RDP and RB substrates twice. The x- and y-axes indicate the normalized turnover rate for the 1^st^ and 2^nd^ reactions. Cluster 1 and cluster 2 show molecules that stayed in the same peaks twice. (D) Changes in the population ratio with different enzyme concentrations (1, 2, and 11 pM) in the single-molecule assay. PTE-revR2 was used for this assay, and its AE activity was measured with the RB substrate. The λ values at 1, 2, and 11 pM were 0.03 ± 0.01, 0.07 ± 0.01, and 0.23 ± 0.01, respectively (mean ± *SD*, n = 2). (E) Metal reconstruction experiment using the cell-free protein synthesis (CFPS) system (mean ± *SD*, n = 2). PTE-revR2 was expressed either in bacteria or using CFPS with or without ZnCl_2_ supplementation, and these were used to measure population changes between peaks. The graphs in the bar plot indicate the activity distribution for each condition.

Next, we hypothesized that the two peaks might represent different oligomeric states with slightly different activities (**Fig. 3A**). To analyze oligomeric states, fast protein liquid chromatography (FPLC) and blue native PAGE were performed with PTE-R0, revR2, and revR12 (**Fig. S10**) (see **Supplementary Methods**). However, despite the significantly higher enzyme concentrations used in these experiments compared to those in the single-molecule kinetic assay, PTE remained entirely in its monomeric form. Moreover, the population ratio in single-molecule kinetic assay did not significantly change with varying enzyme concentrations (**Fig. 3D**), indicating that the bimodal distribution is unlikely to result from a monomer-dimer transition.

Finally, we hypothesized that the differences in activity might result from heterogeneity of the active site metal ions (**Fig. 3A**). Previous work has shown that PTEs expressed *in vivo* incorporate different metal ions into their binuclear active site [40, 41]. To analyze metal heterogeneity, we utilized *in vitro* cell-free protein synthesis (CFPS) to express revR2 with or without excess ZnCl_2_ (see **Supplementary Methods**) [32, 34]. When revR2 was expressed without zinc supplementation, the population ratio was 0.18 ± 0.04 (mean ± *SD*, n = 2) — essentially entirely in peak 1L (**Fig. 3E**). With zinc supplementation, the population ratio increased to 1.50 ± 0.04, which is similar to the ratio observed for bacterial expression supplemented with ZnCl_2_ in a growth media (1.39 ± 0.01) (**Fig. 3E**). This is fully consistent with previous work showing that PTEs incorporate a Fe^2+^-Zn^2+^ active site in the absence of excess Zn^2+^ [41], and that the Zn^2+^- Zn^2+^ active site variant exhibits higher activity [35, 41].

The bimodal distribution appeared to result from the formation of different active site metal ion centers. However, the differences in activity between the two peaks and the variation in their populations are relatively small (**Fig. S7A-B**), compared to the magnitude of the activity change across the evolutionary trajectory (**Fig. 2B**). In other words, the different variants do not exhibit 10^4^-fold higher or lower activity due to large changes in metal ion incorporation. Rather, the differences in activity are related to the specific activity of each variant. Regardless, for the following analysis, we focused on the most active peak 1H, while also performing analyses on the combined peak 1L and peak 1H as a control (see **Supplementary Methods**). Note that the total width of peak 1, evaluated by the coefficient of variation (*CV*_n_), did not differ significantly between bacterial and CFPS expression with or without Zn^2+^ (*CV*_n_ values for CFPS with and without Zn^2+^and for bacterial expression were 19 ± 2%, 21 ± 0.4%, and 17 ± 4% (mean ± *SD*, n = 2), respectively (p-value > 0.1)). Moreover, *CV* in peak 1L and peak 1H exhibited a positive correlation (**Fig. S7D**), indicating that the molecular cause of the bimodal distribution did not significantly affect the width of the activity.

### Evolution of functional heterogeneity in PTE variants

The second type of heterogeneity we observed in the single-molecule data was the width of the activity distribution. To quantify the degree of heterogeneity, we calculated the coefficient of variation (*CV*, %) or relative standard deviation of peak 1H (see **Supplementary Methods**). Most of the enzyme variants exhibited a wider distribution (*CV* = 10-23% in **Fig. 4A**) than the detection limit with our system (*CV*_limit_ = 9.3 ± 1.0%, mean ± *SE*, n = 4), which is calculated from the spontaneous reaction rate of the substrate. Interestingly, we found that the *CV* values gradually changed throughout the evolution, and the change correlated with the functional transition (**Fig. 2B** and **Fig. 4A**). For instance, the *CV*_PTE_ remained around 15% for the first eight rounds of evolution, during which variants exhibited PTE-specific or bifunctional features, before dropping rapidly to around 10% as the specialization for AE activity progressed (**Fig. 4A**). The same trend is observed in reverse evolution, with the *CV*_PTE_ increasing to 15%, as PTE activity was restored. The changes in *CV*_AE_ exhibited a similar trend, with a gradual decrease to 10% at R22, and then an increase during the reverse evolution (**Fig. 4A**). Since mutations accumulated during the evolutionary process, we could observe the trends above in PTE and AE when we plotted the accumulation of mutations and *CV* (**Fig. S11**). Note that the kinetic system itself, such as the binding of enzyme molecules, did not affect *CV* (**Fig. S6B-C** and **Fig. S8D**).

**Figure 4.**
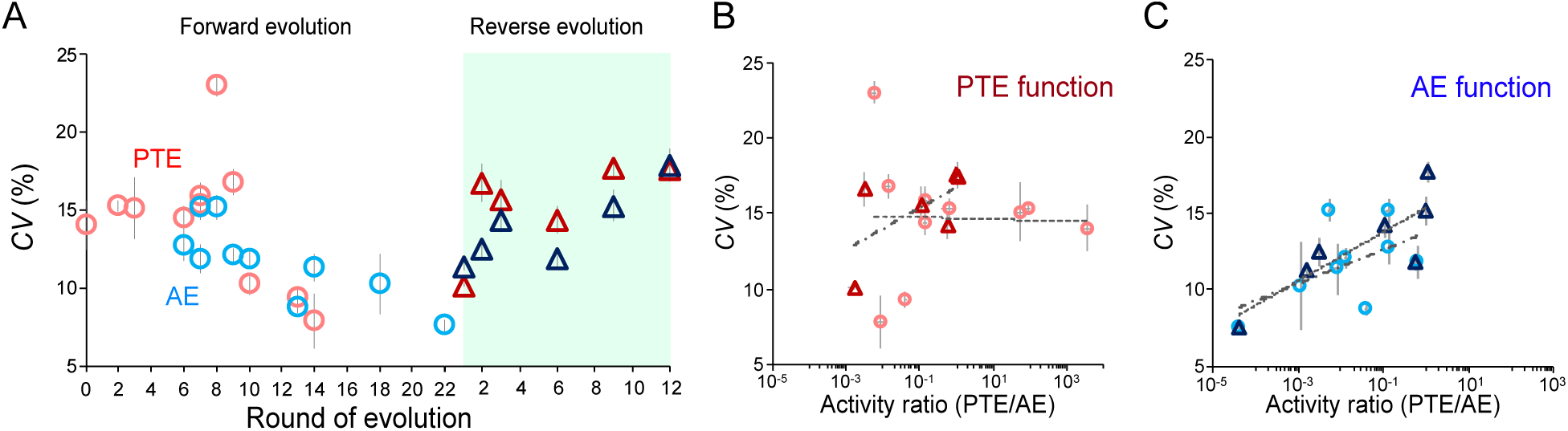
Change of functional substates in enzyme evolution measured by single-molecule kinetic assay. (A) Change of functional substates (*CV*, %) of peak 1H during evolution. *CV*s were measured with RDP (Red circles and triangles) and RB (blue circles and triangles) substrates (mean ± *SE*, **Table S1**). Forward and reverse evolutions are shown as circles and triangles, respectively. (B)-(C) Correlation between *k*_cat_/*K*_M_ ratio of PTE against AE and functional substates. The x-axis was converted to a logarithmic scale. The dashed and dashed-dotted lines show log-linear fittings for the plots of the forward and reverse evolutions: -0.0003 (Pearson’s correlation coefficient = -0.033) and 0.006 (0.38) in (B), and 0.005 (0.62) and 0.007 (0.85) in (C), respectively.

The *CV* showed a clear trend when compared to the activity ratio (*k*_cat_/*K*_M_; PTE/AE): AE-specialized variants (PTE/AE < 10^-1^) exhibited low *CV*, while bifunctional variants (10^-1^ < PTE/AE < 10^1^), as well as PTE-specialized variants (PTE/AE > 10^1^), displayed higher *CV* values (**Fig. 4B-C**). When we expanded this analysis to include peak 1L, the trends and observations remained unchanged (**Fig. S12**). In other words, peak 1L is essentially the same enzyme, having identical structural mechanisms for function albeit with systematically lower activity due to differences in metal ion incorporation (**Fig. S7D**). These changes in the variation of peak 1 were associated with the evolution of PTE, *i.e*., the activity of the enzymes was more heterogeneous for PTE-specific and bifunctional variants, while the activity distribution associated with AE-specialized variants was considerably narrower. These results suggest that there are mechanistic differences between PTE and AE activities that may influence the *CV*.

### Correlated alterations between conformational heterogeneity and functional substates

The single-molecule activity distributions or functional substate (represented by *CV* values) clearly indicate that PTE activity is associated with a broad functional substate, while AE activity is characterized by a narrower substate. We hypothesized that this could reflect two distinct conformational substates that stably exist in terms of their functional substate across evolution, *i.e.*, the conformational substates resulting in the broad activity distributions are “frozen out” in the forward evolution and restored in the reverse evolution. To address this, we assessed the conformational heterogeneity and dynamics of PTEs by analyzing previously obtained crystal structures of eight PTE variants [35–37]. PTE has several regions that are important for local structural dynamics, such as His/Arg254 and loops 4, 5, and 7 (**Fig. 5A**). We previously demonstrated that their dynamics gradually changed across the evolutionary trajectory and mutational accumulation (**Fig. 5B, Fig. S13A,** and **Fig. S14A**) [37]. For example, loop 7 is highly mobile in the PTE-specific and bifunctional variants and undergoes functionally essential conformational dynamics between open (for substrate binding and/or product release) and closed (catalytic) conformations for PTE substrates, but not for AE substrates [4]. Similarly, a key mutation in the forward evolution, His254Arg, showed conformational heterogeneity. Specifically, a bent conformer of Arg254 stabilizes the naphthol-leaving group of AE substrates, while the alternative extended conformer clashes with the substrate. Critically, loop 7 stabilizes the bent Arg254 conformer in the closed state, and the stabilization of the closed conformation of loop 7 (freezing out the open state that is required for PTE) to enrich the sampling of the bent Arg254 conformer is a central aspect of the evolution of AE specificity. In the reverse evolution, the mobility of loop 7, *i.e.,* sampling of the open state, was restored to regain PTE activity. While loop 7 is stabilized in the closed state in the AE-specific variants, loops 4 and 5, on the opposite side of the active site cleft, become more mobile in AE-specific variants. Loops 4 and 5 likely gained mobility to compensate for the rigidification of loop 7, although the magnitude of their conformational fluctuation is much smaller than that of loop 7 (**Fig. 5B, Fig. S13A,** and **Fig. S14A**).

**Figure 5.**
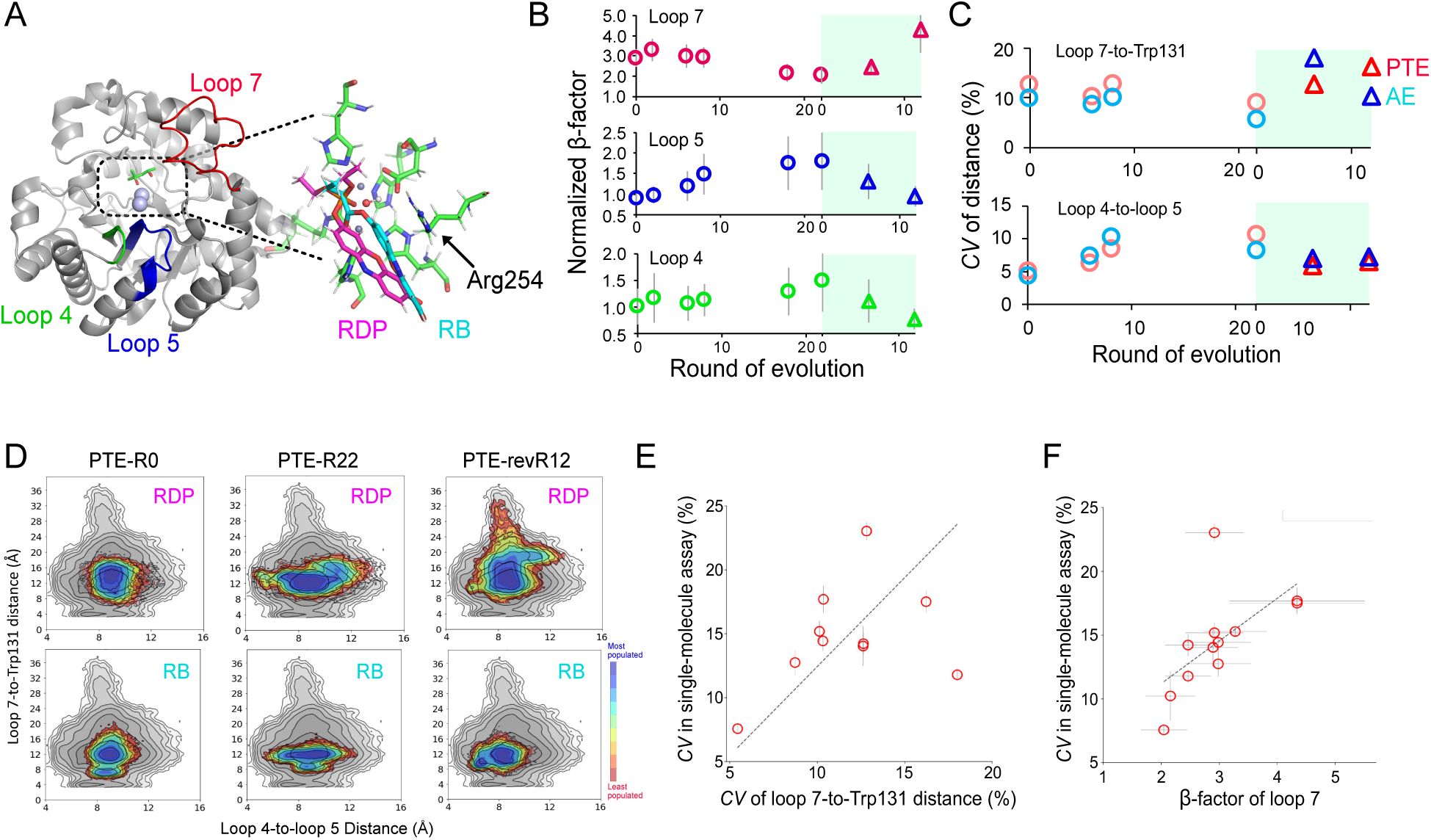
Alteration of local conformational dynamics in enzyme evolution. (A) Functionally important loop structures (loops 4, 5, and 7) for substrate binding. The side chain of Arg254 interacted with the leaving group (resorufin) of both RDP and RB substrates. (B) Change of normalized β-factor in the loop structures. β-factor was analyzed by using crystal structural data of PTE-R0, 2, 6, 8, 18, 22, revR6, and revR12 (mean ± *SD*, n = 2). (C) The change of conformational heterogeneity obtained from MD trajectories. Conformational heterogeneity was analyzed by calculating the coefficient of variation (*CV*, %) in 1D histograms (**Fig. S16A, C**). *CV* was measured with RDP (Red circles and triangles) and RB (blue circles and triangles) substrates. Forward and reverse evolutions were shown as circles and triangles, respectively. (D) Conformational landscape reconstructed considering loops 4, 5, and 7 with RDP and RB substrates. The dynamics of PTE-R0, R22, and revR12 were shown. Blue and red color indicate the degree of the population (blue, more populated) in each state that enzyme sampled. The gray color shows the entire conformational space that the enzyme samples throughout evolution. (E)-(F) Correlation plots of the conformational dynamics (*CV*_loop7_ and β-factor of loop 7) and functional substate (*CV* of peak 1H). The linear fittings of the data points provided fitting slopes; 1.4 (Pearson’s correlation coefficient = 0.4) in (E) and 0.03 (0.6) in (F), respectively.

To gain further insight into the potential correlation between the dynamics of PTEs and functional substates observed at the single-molecule level, we performed molecular dynamics (MD) simulations with RDP and RB substrates, as well as in the absence of a substrate (apo state). To explore PTE conformational dynamics on functionally relevant time scales, we employed Gaussian Accelerated MD (GaMD) to enhance conformational sampling (see **Supplementary Methods**). We first analyzed the β-factors of loops 4, 5, and 7. The β-factors in MD simulation reflected the trends observed in the crystal structure analysis. In the apo state, the β-factor of loop 7 was greater in the variants with higher PTE activity (**Fig. S13B**). Substrate binding significantly reduced the β-factor of loop 7, indicating a shift toward closed conformational states upon ligand binding (**Fig. S13B-D**). For loops 4 and 5, substrate binding did not largely reduce the β-factor. The β-factor increased in the apo state along the AE (forward) trajectory (**Fig. S13B**), which is again consistent with the crystallographic data.

Next, we explored the conformational heterogeneity from the MD analysis by measuring the distance between Ala270 in loop 7 and Trp131, which is located on the opposite side of loop 7 with less fluctuation in MD trajectory (as a proxy for the sampling of the open state), and reconstructed the conformational landscapes (**Fig. 5D**, **Fig. S15**, and **Fig. S16A-B**). In the apo state, R0 exhibited two distinct peaks at 14 and 21 Å, corresponding to catalytic (close) and substrate binding/product release (open) states. In the forward evolution, the population of the open conformation significantly decreased, with the mean distance reduced from 16 ± 4 Å in R0 to 9 ± 4 Å in R22. In the reverse evolution (to improve PTE activity), the sampling of the open state was restored, with the mean distance increasing from R22 (9 ± 4 Å) to revR12 (14 ± 4 Å). For loops 4 and 5, we measured the distance between the central amino acids of loop 4 (Thr173-Val176) and loop 5 (Ala203-Gln206). Unlike loop 7, clear conformational states were not observed in these loops; instead, they appear to exhibit short-range dynamics, rather than fluctuations between well-defined states. Indeed, these loops are much smaller and more conformationally restricted (**Fig. S15** and **Fig. S16C**-**D**).

We then quantitatively analyzed the conformational heterogeneity of the loops in substrate-bound states by calculating the coefficient of variations (*CV*_loop7_ and *CV*_loop4-5_) from the 1D histograms (**Fig. S16**) (see **Supplementary Methods**) and plotting them against the evolutionary round and mutational accumulation (**Fig. 5C** and **Fig. S14B**). The overall trends of *CV* changes in the MD simulation were almost identical to those of crystal structures; *CV*_loop7_ decreased in the forward evolution and was then restored in the reverse evolution. *CV*_loop4-5_ showed trends opposite to those of *CV*_loop7_. In addition, *CV*_loop7_ (12.2 ± 2.5% and 10.4 ± 4% with RDP and RB, mean ± *SD*) was higher than *CV*_loop4-5_ (7.2 ± 2.0% and 7.4 ± 2.0%) in enzyme evolution, indicating large conformational dynamics between open and closed conformal substates in loop 7. To clarify the possible coupling between conformational heterogeneity and functional substate, we analyzed the correlation between *CV*_loop7_ or β-factor in the crystal structure and *CV* in single-molecule kinetic assays. Interestingly, both conformational and functional heterogeneity exhibited positive correlations (**Fig. 5E-F**). These results indicate that *CV* in single-molecule kinetic assays reflects the dynamics of loop 7 and/or its related global structural dynamics and that the reduction in *CV* observed during the forward evolution of AE function is caused by the ’freezing out’ of the associated loop 7 heterogeneity. Consequently, reverse evolution leads to the recovery of *CV* through the reintroduction of loop 7 heterogeneity. Finally, we assessed the alteration of global structural dynamics involved in loop 7 during enzyme evolution using shortest path maps (SPMs) derived from MD simulation data [20, 21, 42, 43]. SPM captures pairs of residues that exhibited correlated dynamics during MD simulation. We highlighted these correlated networks on the 3D structure, with spheres representing individual residues and thicker edges indicating correlated residue pairs, weighted based on their correlation values (**Fig. 6A** and **Fig. S17A**) [20, 21, 42, 43]. In forward evolution, the number of edges increased at substrate-bound states, which would minimize conformational dynamics and facilitate the sampling of more closed states for the AE function (**Fig. S17A** and **Fig. S18A**). Indeed, the edges between Arg254 and Phe/Leu271, which stabilize closed conformational states essential for AE function, were observed in both bifunctional and AE-specialized enzymes (**Fig. 6B** and **Fig. S17B**). In the reverse evolution of the RDP binding state, the total number of edges decreased, and edges between Arg254 and Leu271 were lost, leading to the recovery of loop7 dynamics. In other words, since the extended networks constrain the dynamics of loop 7, which is crucial for PTE activity, the number of edges was inversely correlated with the *CV*_PTE_ measured in the single-molecule kinetic assay (**Fig. S18B**).

**Figure 6.**
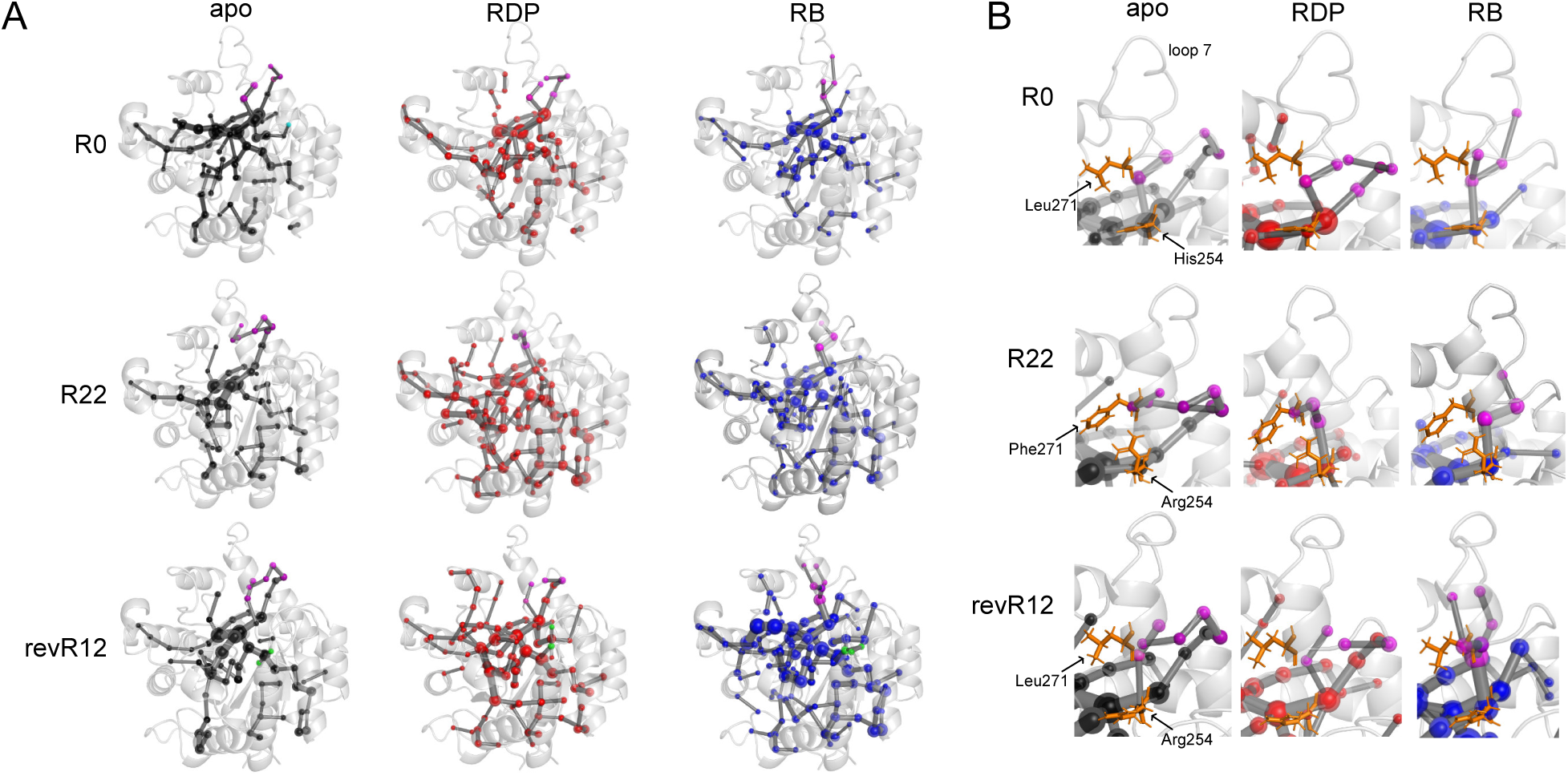
Shortest path maps (SPMs) of PTE variants. (A) Alteration of SPMs in PTE-R0, R22, and revR12 were analyzed in the absence of a substrate (apo-state, black sphere) and with RDP (red sphere) and AE (blue sphere) substrates. Spheres and edges represent individual residues and correlated residue pairs, respectively. SPMs in loops 5 and 7 are shown as cyan and magenta spheres, respectively. (B) Alteration of SPMs on loop 7.

## Conclusion

We performed a single-molecule enzyme kinetic assay using a microreactor array device to measure the heterogeneous functional substates of PTE and its evolutionary trajectories. Changes in functional substates, coupled with alterations in local structural dynamics critical for PTE function, revealed their correlation for the first time. Our results suggest that coordinated changes across different dynamic hierarchies are essential for enzymes to acquire and refine new functions throughout evolution. Expanding the analysis to a broader range of enzymes will enhance our understanding of how functional substates relate to structural and biophysical properties, shedding light on the roles of heterogeneous enzyme ensembles in evolution. A deeper understanding of functional substates could further drive advances in enzyme engineering and rational protein design.

## Acknowledgment

The authors are grateful to members of the Tokuriki lab for helpful discussions and technical support. M.S. is supported by the Japan Society for the Promotion of Science (JSPS) Overseas Research Fellowship. F.F. received the support of the Spanish MICIU (Ministerio de Ciencia, Innovación y Universidades) (Projects PID2022-141676NB-I00 and TED2021-130173B-C42 and fellowship RYC2020-029552-I) and Generalitat de Catalunya for the consolidated group TCBioSys (SGR 2021-00487). E.N. is supported by the Grant-in-Aid for Transformative Research Areas (A) from the Japan Society for the Promotion of Science (22H05418 and 24H01129). S.O. received the support of the Generalitat de Catalunya for the consolidated group TCBioSys (SGR 2021 00487), Spanish MICIN for grant projects PID2021-129034NB-I00 and PDC2022-133950-I00. S.O. is grateful to the funding from the European Research Council (ERC) under the European Union’s Horizon 2020 research and innovation program (ERC-2015-StG-679001, ERC-2022-POC-101112805, ERC-2022-CoG-101088032, and ERC-2023-POC-101158166). N.T. is supported by the Natural Sciences and Engineering Research Council of Canada (NSERC) / Discovery Grants Program (RGPIN-2023-05135). N.T. and S.O. are supported by the Human Frontier Science Program (HFSP) Research Grant (RGP0054/2020).

## Author contributions

M.S. conceived the project, designed experiments, performed experiments, analyzed results, and wrote the manuscript; R.M. and F.F. designed experiments, performed experiments, analyzed results, and wrote the manuscript; C.J.J. conceived the project and wrote the manuscript; E.N. designed experiments, performed experiments, analyzed results, and wrote the manuscript; S.O. designed experiments, performed experiments, analyzed results, and wrote the manuscript; and N.T. conceived the project, designed experiments and wrote the manuscript.

