## Supplementary information for "Single-Molecule Kinetic Exploration of Functional Substates in an Evolving Phosphotriesterase"

**Supplementary Information content:**


**Figure S1** $k_{cat}/K_M$ of purified wild-type PTE-R0 and its variants measured with paraoxon and 2NH substrates

**Figure S2** Microreactor array device for single-molecule kinetic assay

**Figure S3** Evaluation of cross-reactor diffusion of resorufin dye

**Figure S4** Distribution of catalytic turnover rates measured in a single-molecule kinetic assay

**Figure S5** Linear Correlation between fluorescence intensity and resorufin dye concentration

**Figure S6** Effect of single-molecule assay on enzyme function

**Figure S7** Analysis of bimodal distribution

**Figure S8** Assessment of the reaction solution exchange experiment

**Figure S9** Typical scatter plots from the reaction solution exchange experiments

**Figure S10** Characterization of the monomer-dimer transition with PTE-R0, revR2, and revR12

**Figure S11** Correlation between mutational accumulation and functional substates (*CV* in peak 1H) of PTE and AE functions.

**Figure S12** Analysis of the total width of bimodal distribution in enzyme evolution

**Figure S13** Main chain $\beta$-factor of loops 4, 5, and 7

**Figure S14** Correlation between mutational accumulation and conformational dynamics

**Figure S15** 2D histograms of conformational dynamics in functionally important loop structures

**Figure S16** 1D histograms of conformational dynamics in functionally important loop structures

**Figure S17** Shortest path maps (SPMs) of PTEs

**Figure S18** The change of SPMs in PTE and AE activities

**Table S1** Kinetic assay data of PTEs

**Table S2** Reaction solution exchange experiment data

**Supplementary Methods**

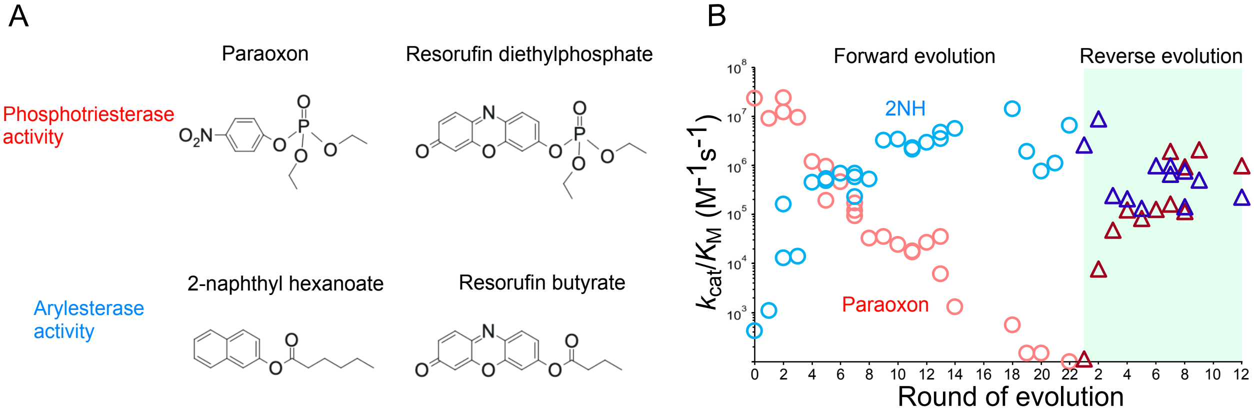


**Figure S1. $k_{cat}/K_M$ of purified wild-type PTE-R0 and its variants measured with paraoxon and 2NH substrates**. (A) Chemical structures of paraoxon, 2-naphthyl hexanoate (2NH), resorufin diethylphosphate (RDP), and resorufin butyrate (RB) substrates. (B) $k_{cat}/K_M$ measured with paraoxon (red) and 2NH (blue) substrates. Forward and reverse evolutions are shown as circles and triangles, respectively. The y-axis was converted to a logarithmic scale. Data are cited from measurements taken in a previous experiment [36].

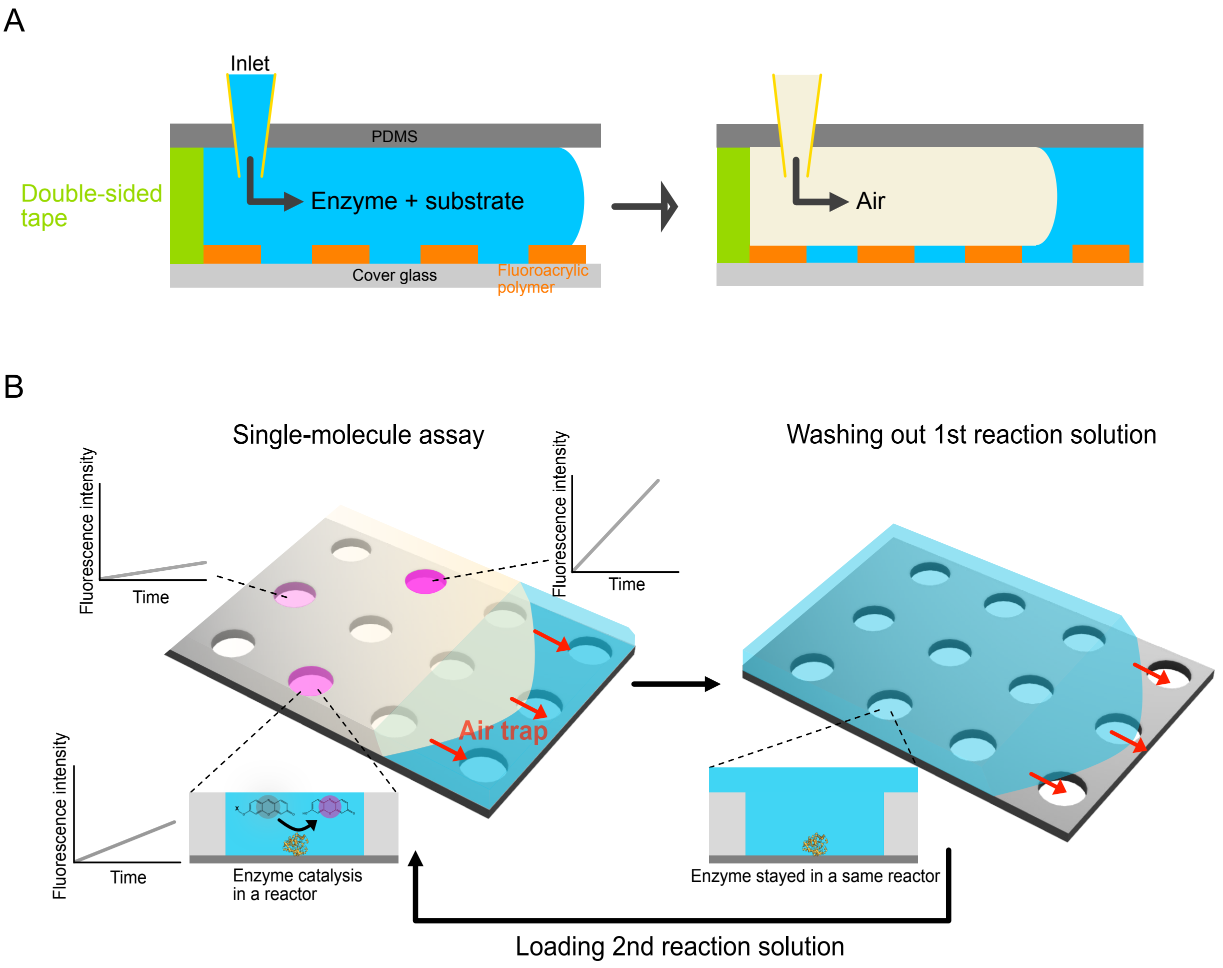


**Figure S2. Microreactor array device for a single-molecule kinetic assay.** (A) The microreactor array was formed on a cover glass using photolithography techniques. Double-sided tape was cut into the shape of a flow channel and applied to the device. The top surface of the device was affixed with PDMS having inlet and outlet holes for injecting solution with a pipette. For a single-molecule assay, a reaction solution containing enzyme and substrate was injected from the inlet. Then, air was injected from the same inlet to remove excess solution and seal the reactors. (B) Reaction solution exchange experiment with the microreactor array device. After measuring 1st reaction, the fresh buffer without substrate was injected, and reactants in the reaction were removed from the reactors. Then, the fresh reaction buffer having substrate was injected to measure 2nd reaction with an identical enzyme molecule.

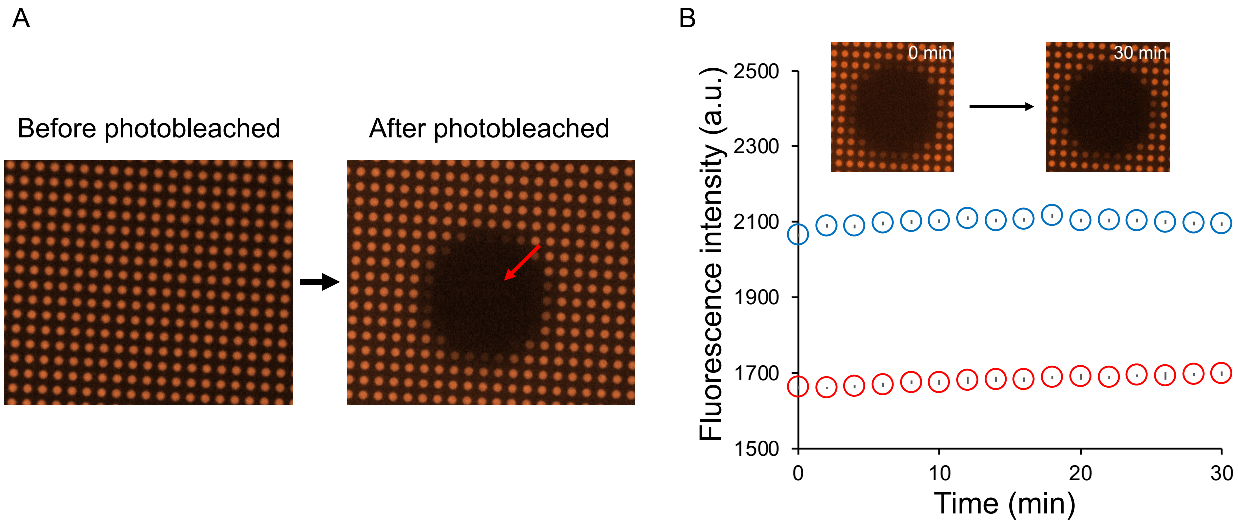


**Figure S3. Evaluating cross-reactor diffusion of resorufin dye.** (A) Resorufin dye was encapsulated in the reactors, and a limited area was photobleached by irradiating strong excitation light (red arrow). (B) Time-course change of fluorescence intensity in photobleached (red circles) and surrounding (blue circles) reactors (mean ± *SD*, n = 6 reactors). Microscope images were taken at 2-minute intervals for 30 minutes.

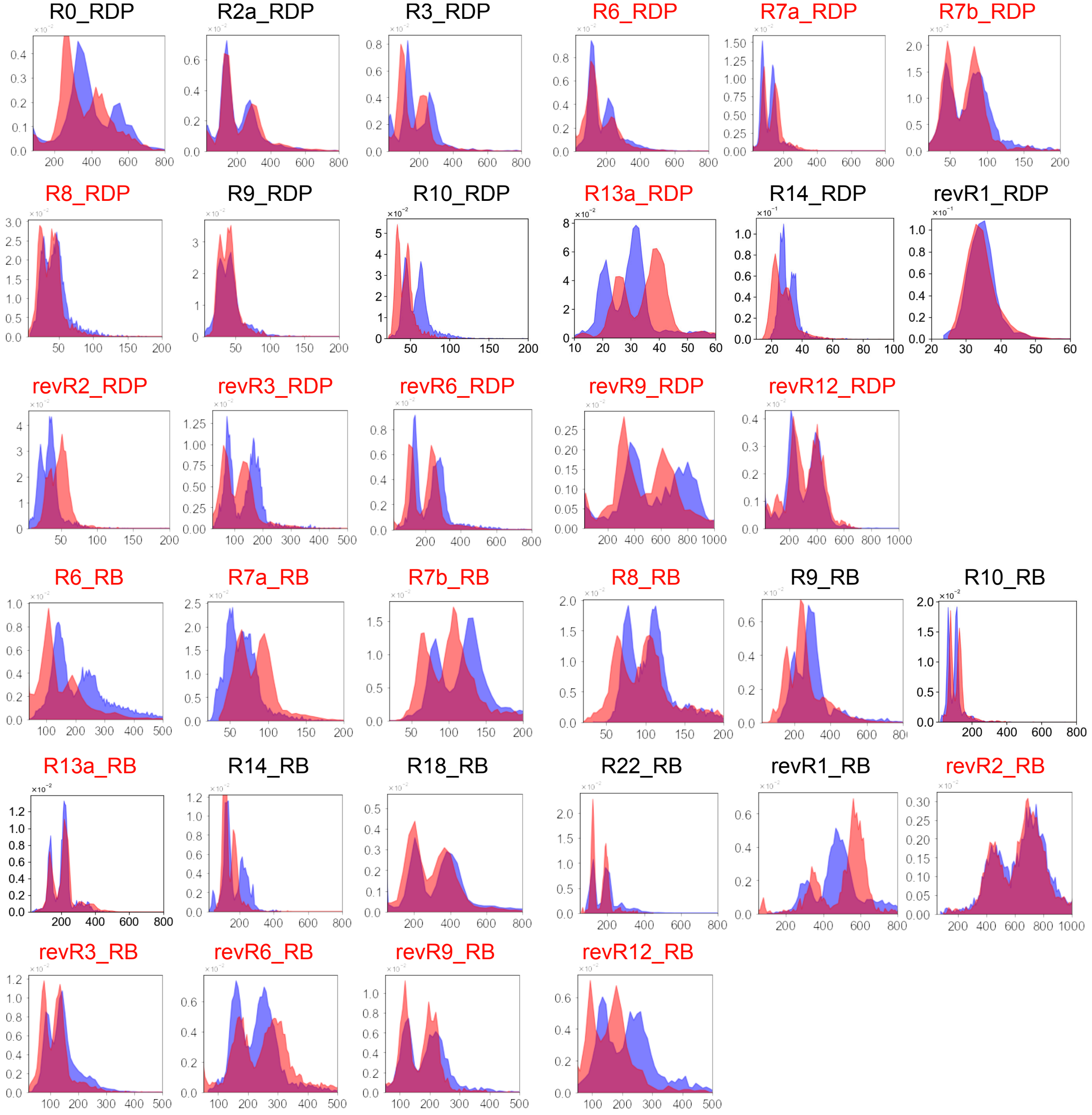


**Figure S4. Distribution of catalytic turnover rates measured in a single-molecule kinetic assay.** The x- and y-axes exhibit turnover rate ($s^{-1}$) and the population percentage of enzyme molecules. The red and blue bins show the distribution of two independent measurements. The total number of enzyme molecules was over 700 in each assay. The identifiers of the mutants used in the reaction solution exchange experiment (**Fig. 3**) are shown in red.

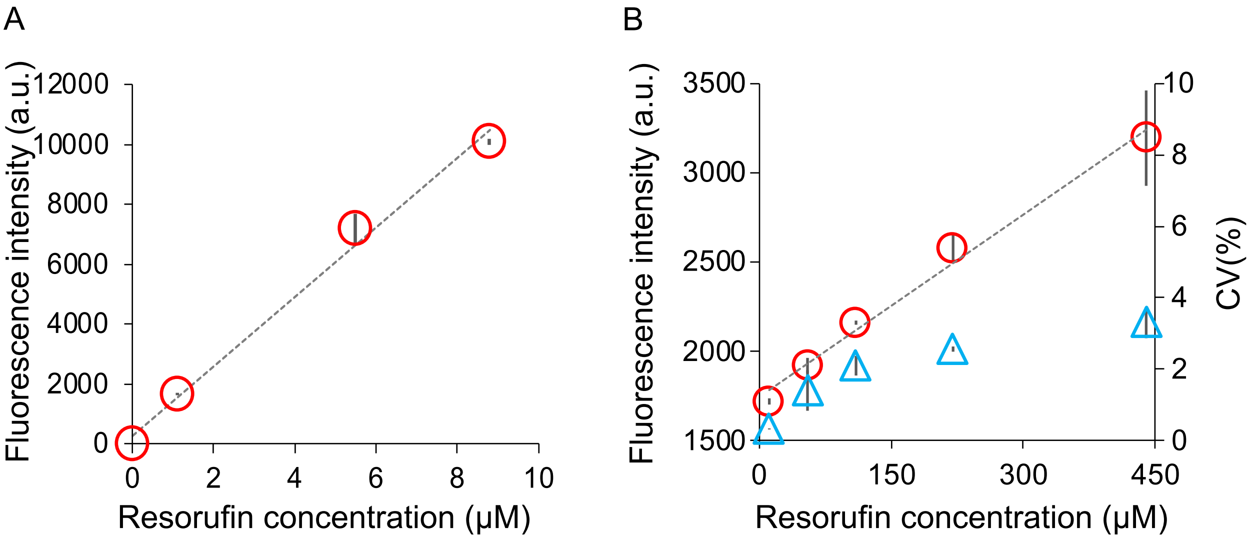


**Figure S5. Linear Correlation between fluorescence intensity and resorufin dye concentration.** (A) Calibration curve measured in a 384-well plate (mean ± *SD*, n = 3). The dashed line shows a linear fit for the plot (slope = 1,153, $R^2$ = 0.99). (B) Calibration curve in a microreactor array device measured with two different devices (mean ± *SD*, n = 3) (red circles). The coefficient of variation of the distribution of the fluorescence intensity was calculated as an inter-reactor error ($CV_{reso}$) (blue rectangles). The dashed line shows a linear fit for the plot (slope = 3.4, $R^2$ = 0.99).

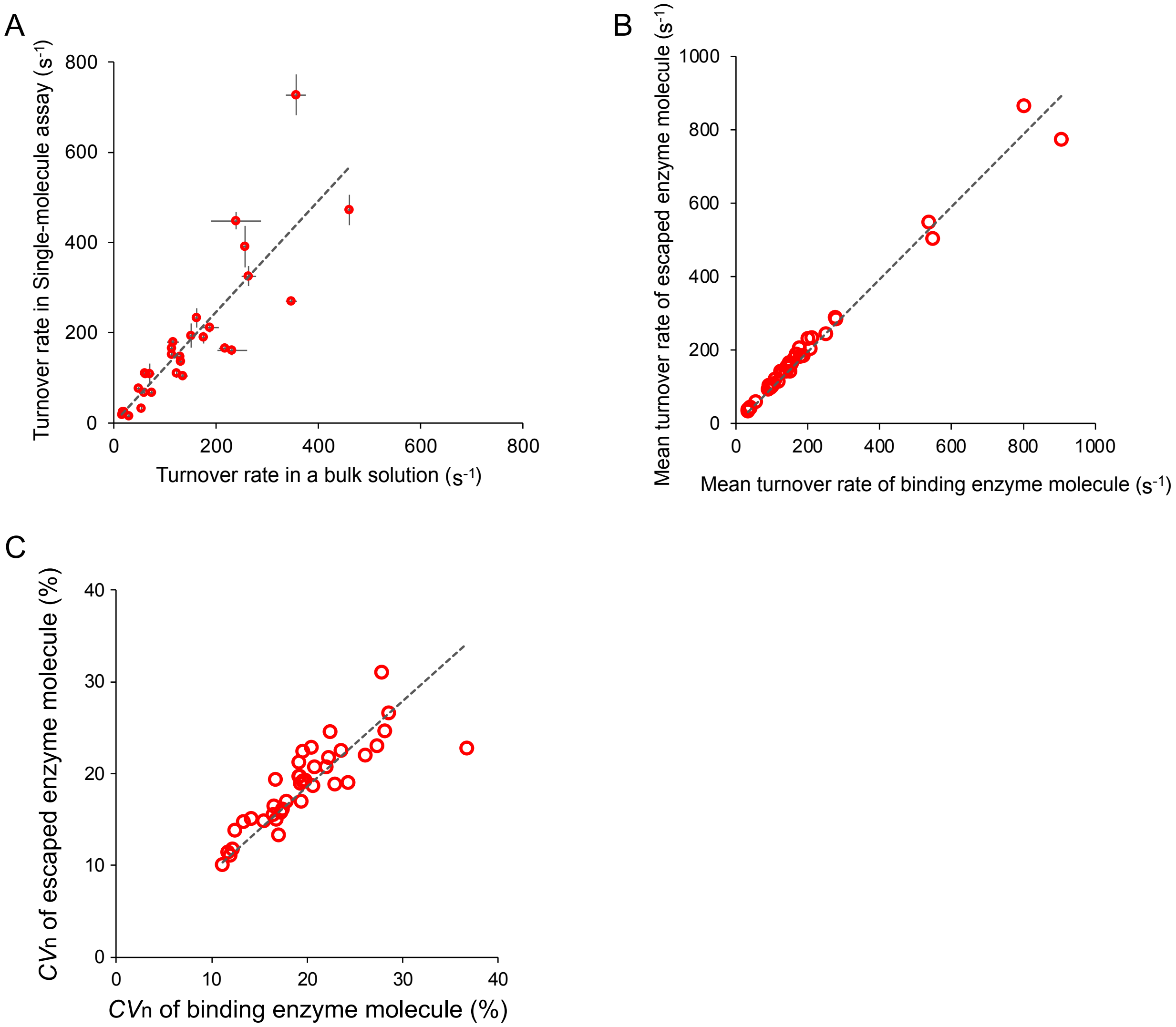


**Figure S6. Effect of single-molecule assay on enzyme function.** (A) Correlation between the catalytic turnover rates measured in bulk ensemble and single-molecule kinetic assays (mean ± *SE*, **Table S1**). The dashed line shows a linear fit for the plot (slope = 1.2, $R^2$ = 0.72). (B), (C) Correlation of mean turnover rate and $CV_n$ between binding and escaped enzyme molecules. Binding and escaped enzyme molecules were identified from the reaction solution exchange experiments ((1st, 2nd) = (PTE, PTE) and (AE, AE) in **Fig. S9**). From the distribution of turnover rate in binding and escaped molecules, the mean runover rate and $CV_n$ were calculated. The dashed lines show linear fits for the plots (slopes = 0.94, $R^2$ = 0.98 and slope = 0.80, $R^2$ = 0.69, respectively).

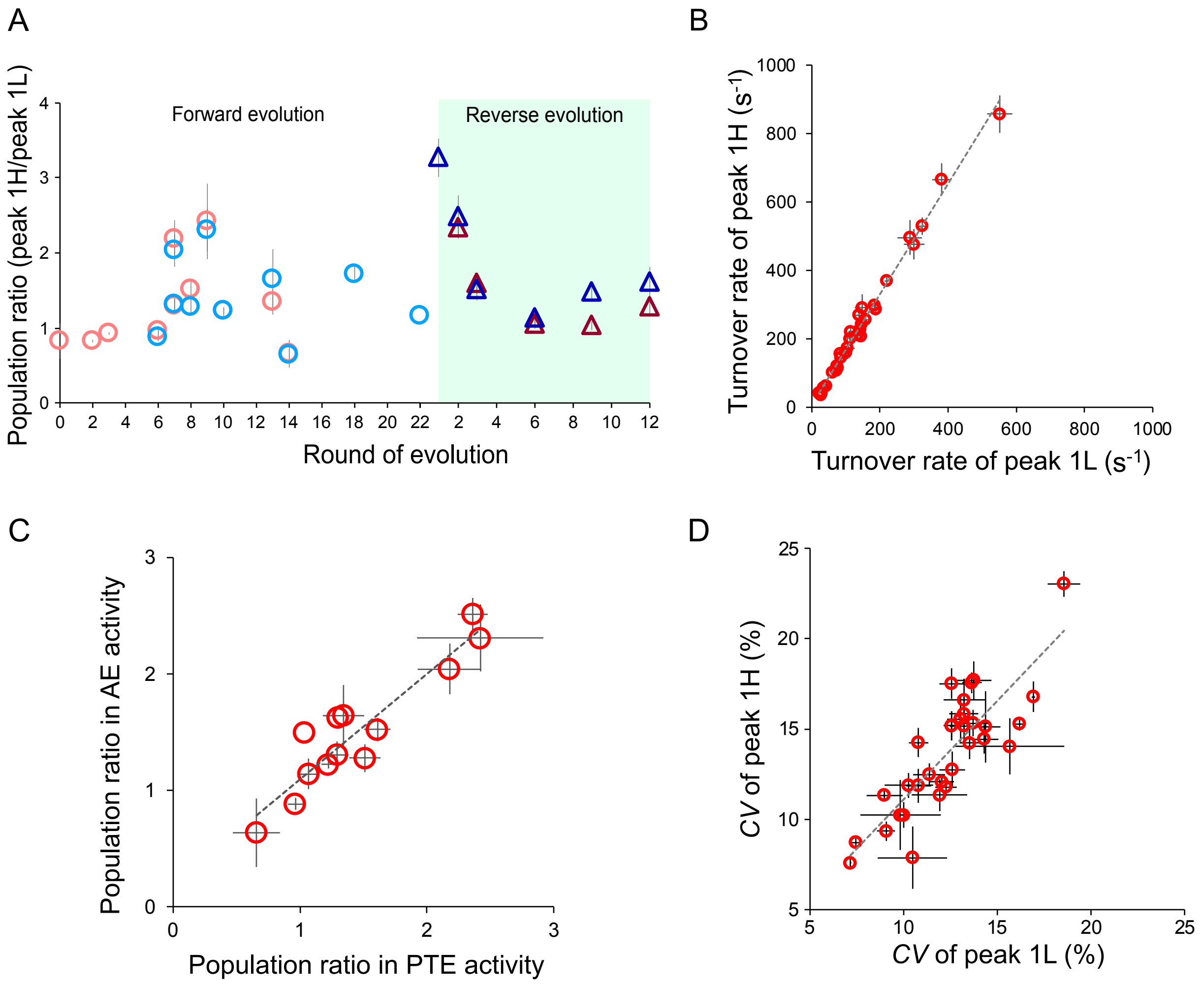


**Figure S7. Analysis of bimodal distribution.** (A) Changes in the population ratio between peak 1H and peak 1L throughout evolution (mean ± *SE*, **Table S1**). Population ratios of peak 1H against peak 1L in PTE and AE functions are shown by red and blue circles or triangles. Forward and reverse evolutions are represented as circles and triangles, respectively. (B) Differences in turnover rate between peak 1H and peak 1L (mean ± *SE*, **Table S1**). The dashed line shows a linear fit for the plot (slope = 1.6, $R^2$ = 0.99). (C) Differences in population ratio in PTE and AE activities (mean ± *SE*, **Table S1**). The plot shows the population ratio for the mutants for which both activities could be measured. The dashed line shows a linear fit for the plot (slope was 0.9, $R^2$ = 0.86). (D) Differences in *CV* between peak1H and peak1L (mean ± *SE*, **Table S1**). The dashed line shows a linear fit for the plot (slope = 1.1, $R^2$ = 0.71).

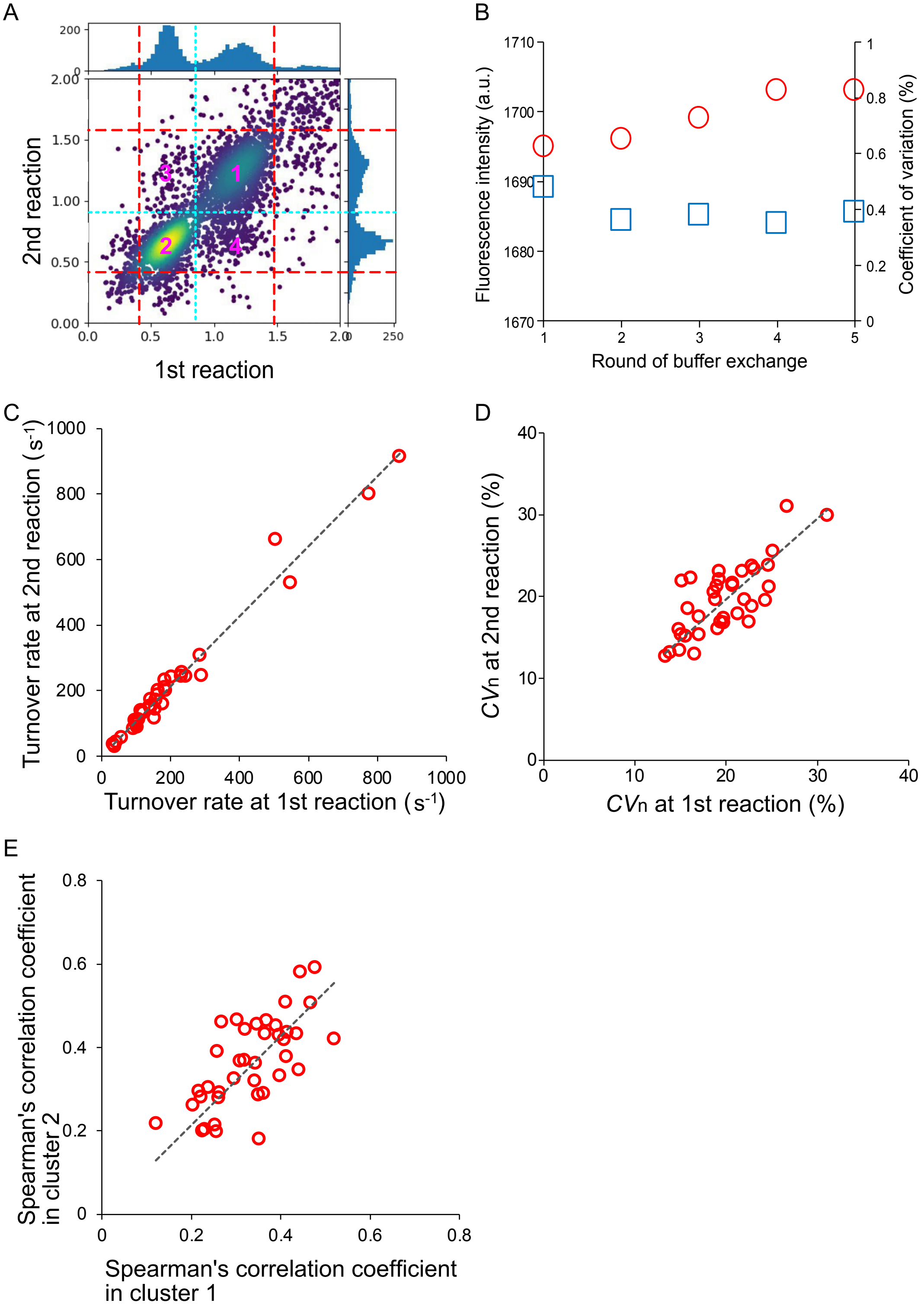


**Figure S8. Assessment of the reaction solution exchange experiment.** (A) Typical data from the reaction solution exchange experiment. Four clusters were observed, and clusters 1 and 2 exhibited enzyme molecules that remained at the same peak in the 2$^{nd}$ reaction. Enzyme molecules in clusters 3 and 4 showed interconversion between the peaks. (B) Four rounds of reaction solution exchange experiments. 10 μM of resorufin dye was successively trapped in the microreactors, and the distribution of the fluorescence intensity was measured. The mean fluorescence intensity and *CV* of the distribution were calculated with over 10,000 reactors. (C), (D) Correlation of mean turnover rate and $CV_n$ between 1st and 2nd reactions in reaction buffer exchange experiment. Data include solution exchange experiments with the same reaction performed twice (($1^{st}$, $2^{nd}$) = (PTE, PTE) and (AE, AE)). The dashed lines show linear fits for the plots (slopes were 1.1, $R^2$ = 0.99 and 0.99, $R^2$ = 0.98, respectively). (E) Positive correlation of Spearman's correlation coefficients between cluster 1 and cluster 2. The dashed line shows linear fits for the plot (slope = 1.1, $R^2$ = 0.95).

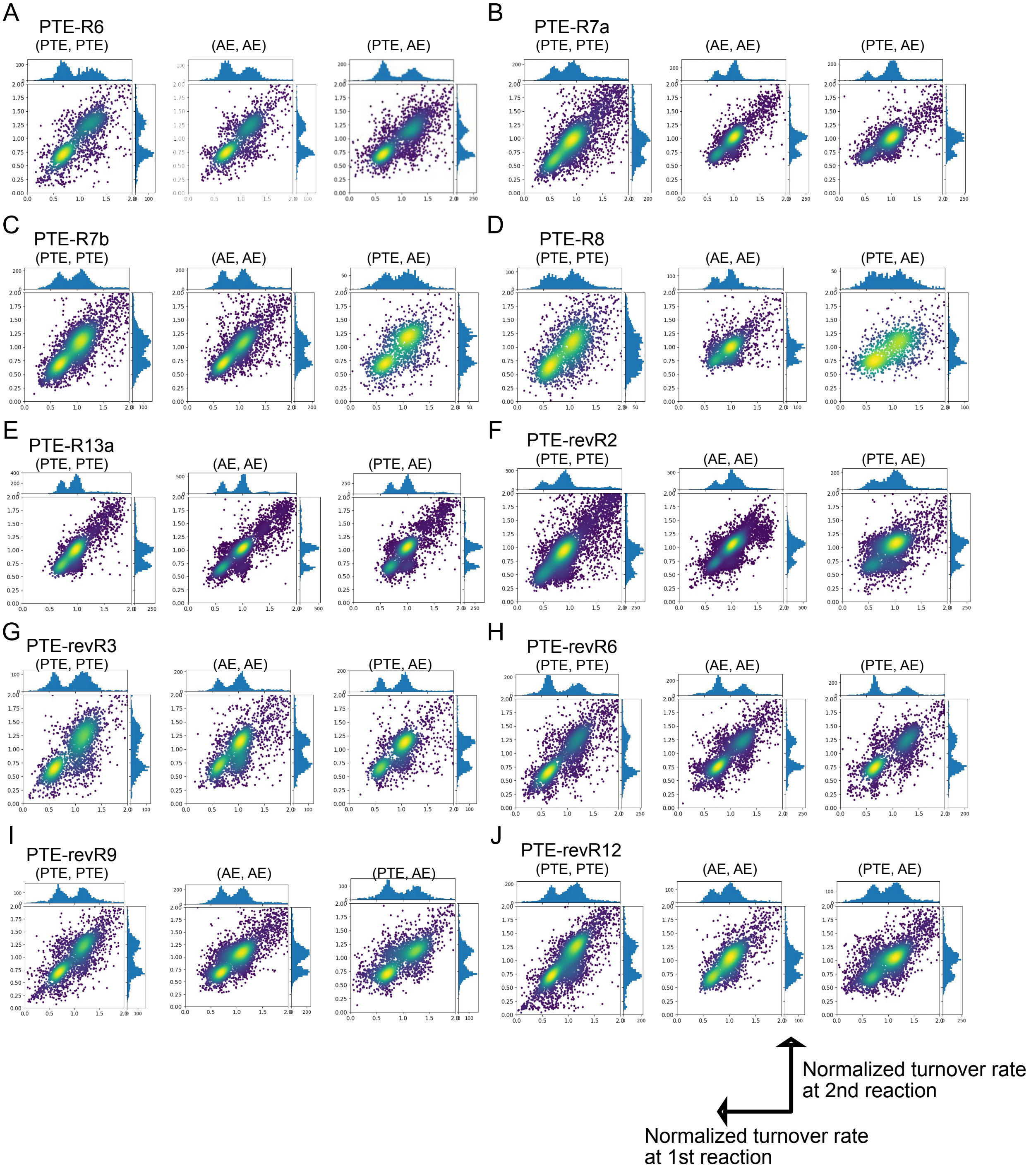


**Figure S9. Typical scatter plots from the reaction solution exchange experiments.** ($1^{st}$, $2^{nd}$) = (PTE, PTE), (AE, AE), (PTE, AE) from left to right.

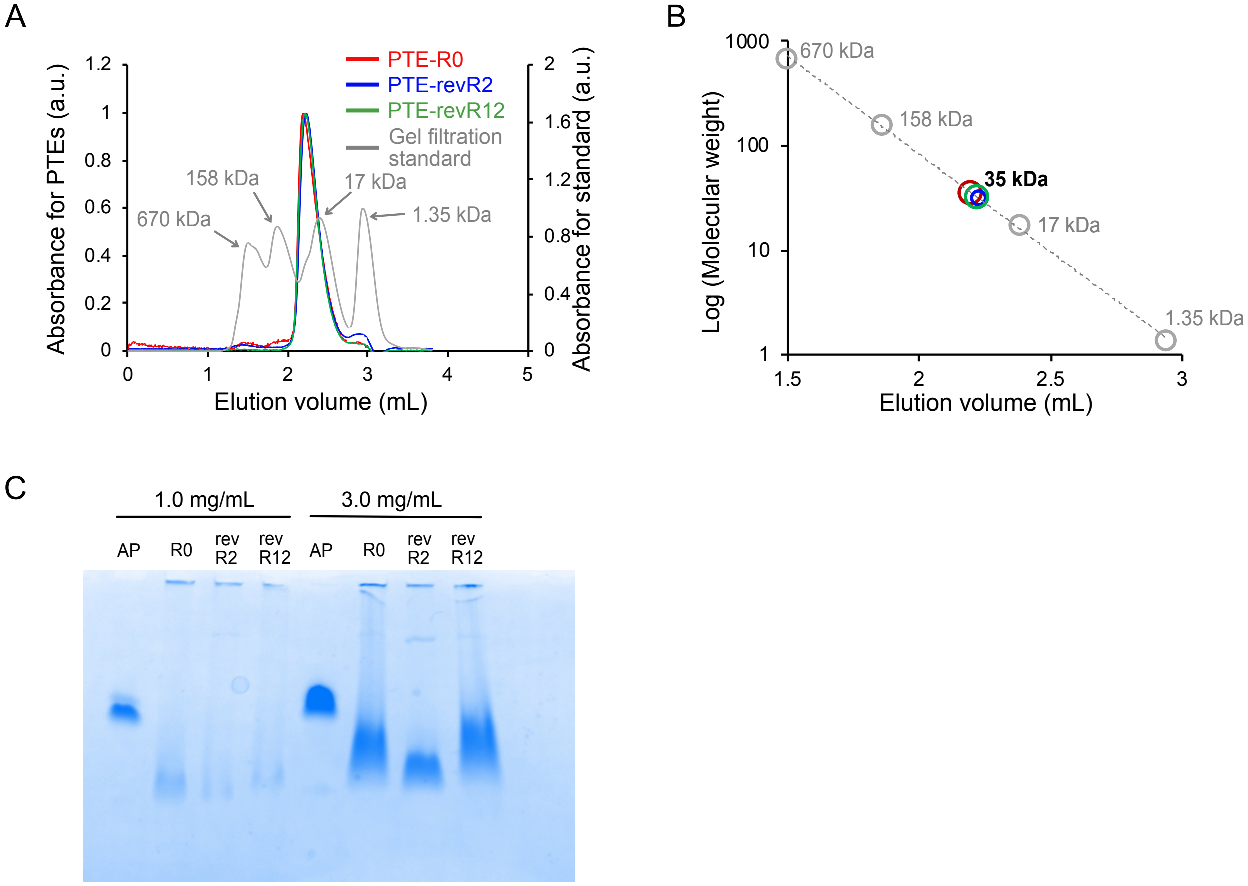


**Figure S10. Characterization of the monomer-dimer transition with PTE-R0, revR2, and revR12.** (A) Size exclusion fast protein liquid chromatography (FPLC) of PTEs and gel filtration standard. The x- and y-axes show the elution volume (mL) and UV absorbance for PTE and the standard. (B) Molecular weight determination from the size exclusion FPLC data. Red, blue, and green circles indicate the molecular weight of R0, revR2, and revR12, respectively. From the linear fitting of the standard, the molecular weight of R0, revR2, and revR12 was estimated as 35.5, 30.9, and 32.3 (kDa), respectively, which are in agreement with the molecular weight of the monomeric form. (C) Blue-native PAGE of PTEs was performed with different concentrations (1.0 and 3.0 mg/mL). *E.coli* alkaline phosphatase (AP, dimer, 86 kDa) was prepared as a positive control. Compared with dimeric AP, PTEs exhibited broad smears at high concentrations, indicating dimer formation at higher protein concentrations.

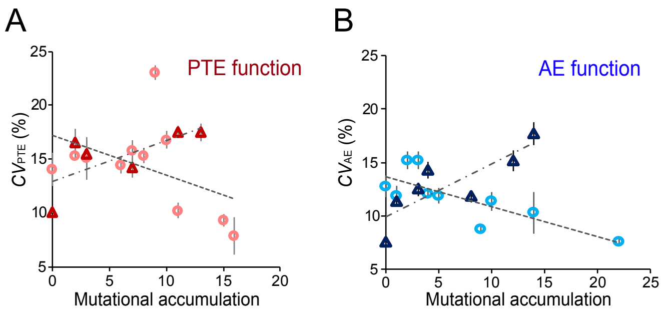


**Figure S11. Correlation between mutational accumulation and functional substates (*CV* in peak 1H) of PTE and AE functions.** (A), (B) The accumulation of mutations for PTE function (A) was calculated from PTE-R0 to R14 (forward evolution, red circles) and from revR1 to revR12 (reverse evolution, blue triangles) (mean ± *SE*, **Table S1**). Accumulation of mutation for AE function (B) was calculated from R6 to R22 (forward evolution, blue circles) and from revR1 to revR12 (reverse evolution, blue triangles) (mean ± *SE*, **Table S1**). The dashed and dashed-dotted lines show linear fittings for the plots of the forward and reverse evolutions; -0.004 ($R^2$ = 0.20, Pearson's correlation coefficient = -0.45) and 0.004 (0.48, 0.69) in (A), and -0.003 (0.63, -0.80) and 0.005 (0.70, 0.84) in (B), respectively.

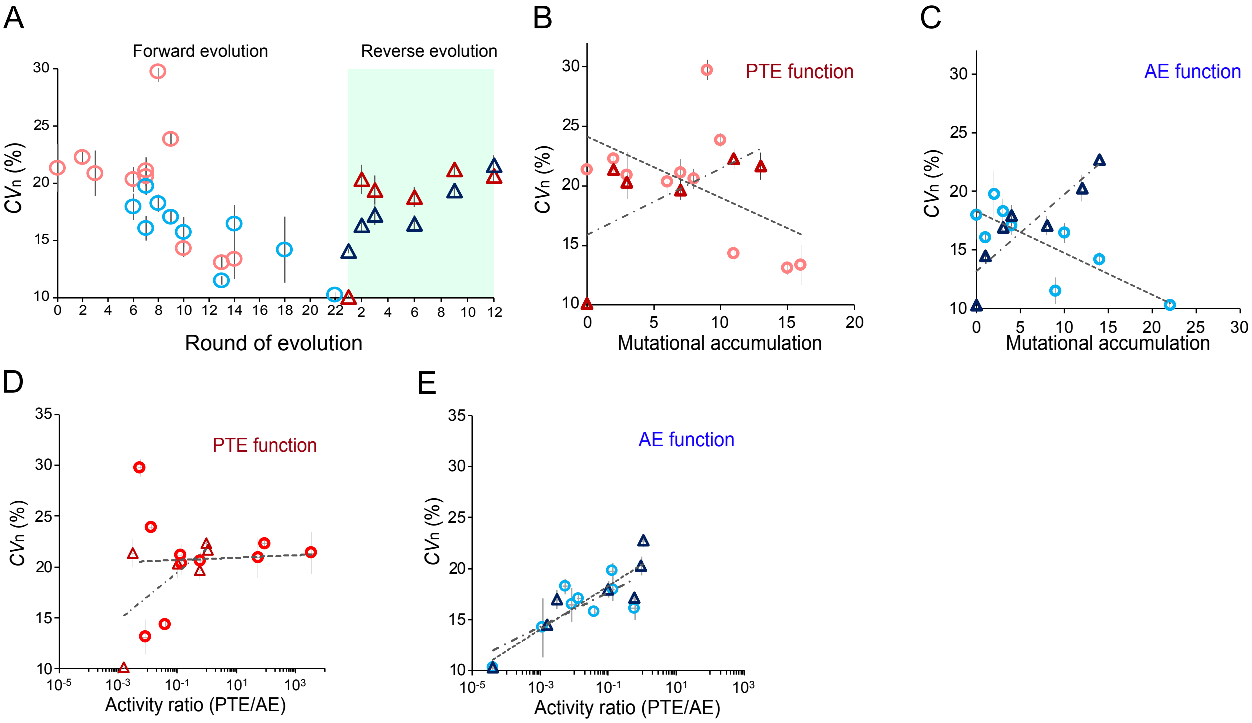


**Figure S12. Analysis of the total width of bimodal distribution in enzyme evolution.** (A) Change of the total width of bimodal distribution ($CV_n$, %) throughout evolution. $CV_n$ of PTEs was measured with RDP (Red circles and triangles) and RB (blue circles and triangles) substrates (mean ± *SE*, **Table S1**). Forward and reverse evolutions are shown as circles and triangles, respectively. (B), (C) Correlation between mutational accumulation and functional substates of PTE and AE functions. Accumulation of mutations for the PTE function was calculated from PTE-R0 to R14 (forward evolution, red circles) and from revR1 to revR12 (reverse evolution, red triangles) (mean ± *SE*, **Table S1**). Accumulation of mutations for the AE function was calculated from R6 to R22 (forward evolution, blue circles) and from revR1 to revR12 (reverse evolution, blue triangles). The dashed and dashed-dotted lines show linear fittings for the plots of the forward and reverse evolutions; -0.004 ($R^2$ = 0.20) and 0.004 (0.48) in (B), and -0.003 (0.63) and 0.005 (0.70) in (C), respectively. (D), (E) Correlation between $k_{cat}/K_M$ ratio of PTE against AE and functional substates (mean ± *SE*, **Table S1**). The x-axis was converted to a logarithmic scale. The dashed and dashed-dotted lines show log-linear fittings for the plots of the forward and reverse evolutions; 0.0003 ($R^2$ = 0.001) and 0.006 (0.38) in (D), and 0.005 ($R^2$ = 0.29) and 0.007 (0.72) in (E), respectively.

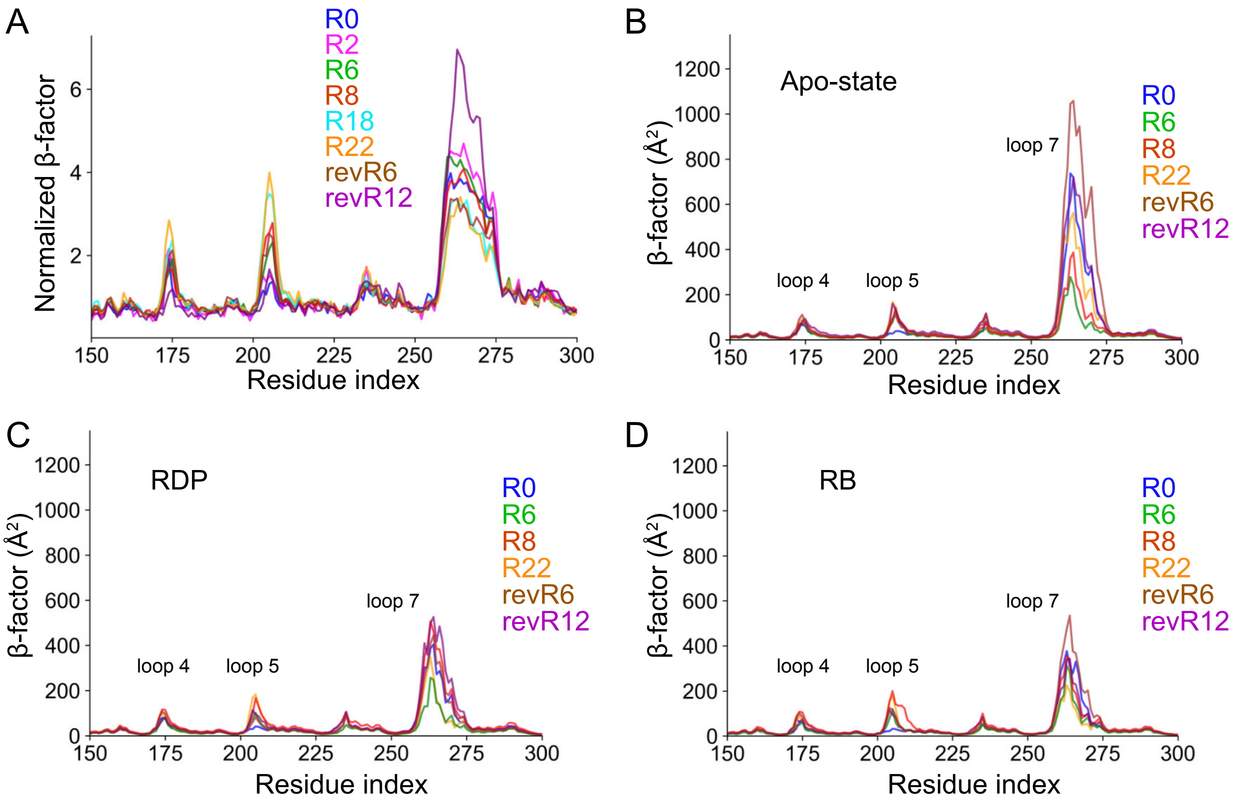


**Figure S13. Main chain β-factor of loops 4, 5, and 7.** (A) The β-factor was analyzed in the crystal structures of PTEs, with the mean β-factor calculated between subunits. (B)-(D) The β-factor was examined in the MD simulations, comparing apo state (substrate-free, B) and substrates-bound states with RDP (C) and RB (D).

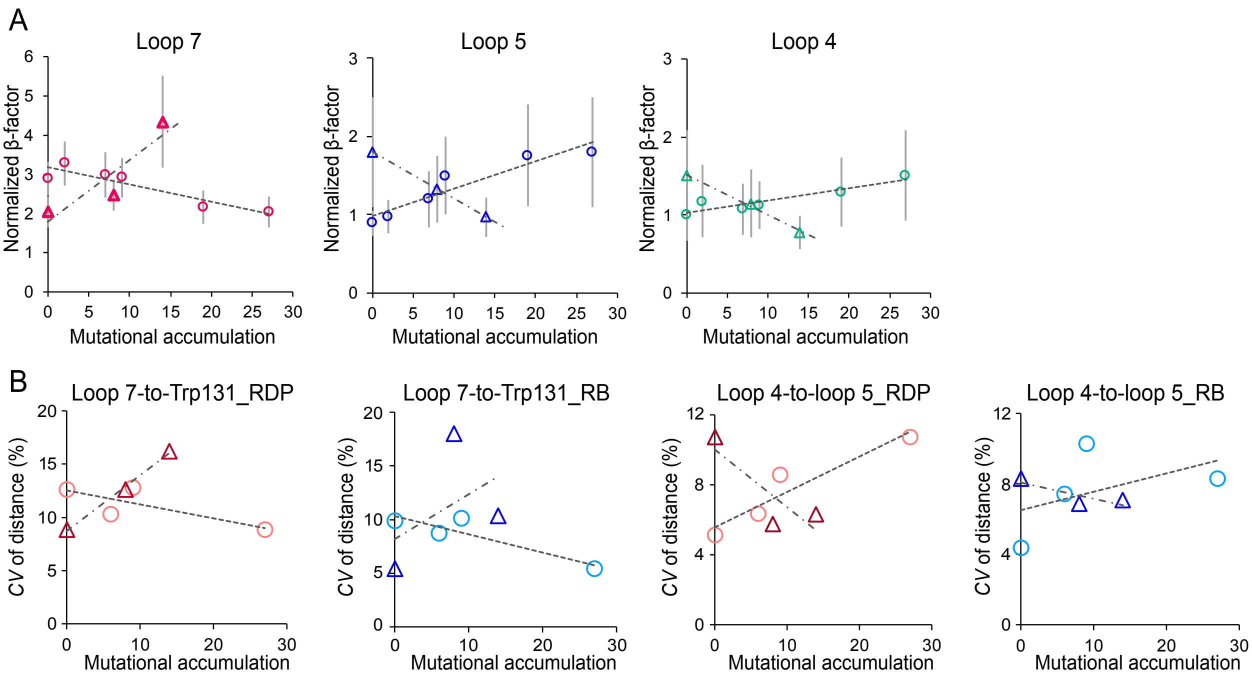


**Figure S14. Correlation between mutational accumulation and conformational dynamics.** (A) Correlations between β-factors of loops and mutational accumulation in the crystal structure data. Accumulation of mutations was calculated from PTE-R0- to R22 (empty circles) and from revR0 (R22) to revR12 (empty triangles). The dashed and dashed-dotted lines show linear fittings for the plots of the forward and reverse evolutions; -0.04 ($R^2$ = 0.85) and 0.16 (0.82) in loop 7, 0.04 (0.89) and -0.06 (0.99) in loop 5, and 0.02 (0.85) and -0.05 (0.99) in loop 4, respectively (mean ± *SD*, n = 2). (B) Correlations between the *CV* of distance in loops and mutational accumulation in MD simulation data. The dashed and dashed-dotted lines show linear fittings for the plots of the forward and reverse evolutions; -0.001 ($R^2$ = 0.61) and 0.005 (0.99) in loop 7-to-Trp131 with RDP, 0.002 (0.83) and 0.004 (0.21) in loop 7-to-Trp131 with RB, 0.002 (0.90) and -0.003 (0.73) in loop 4-to-loop 5 with RDP, 0.001 (0.24) and -0.001 (0.70) in loop 4-to-loop 5 with RB, respectively.

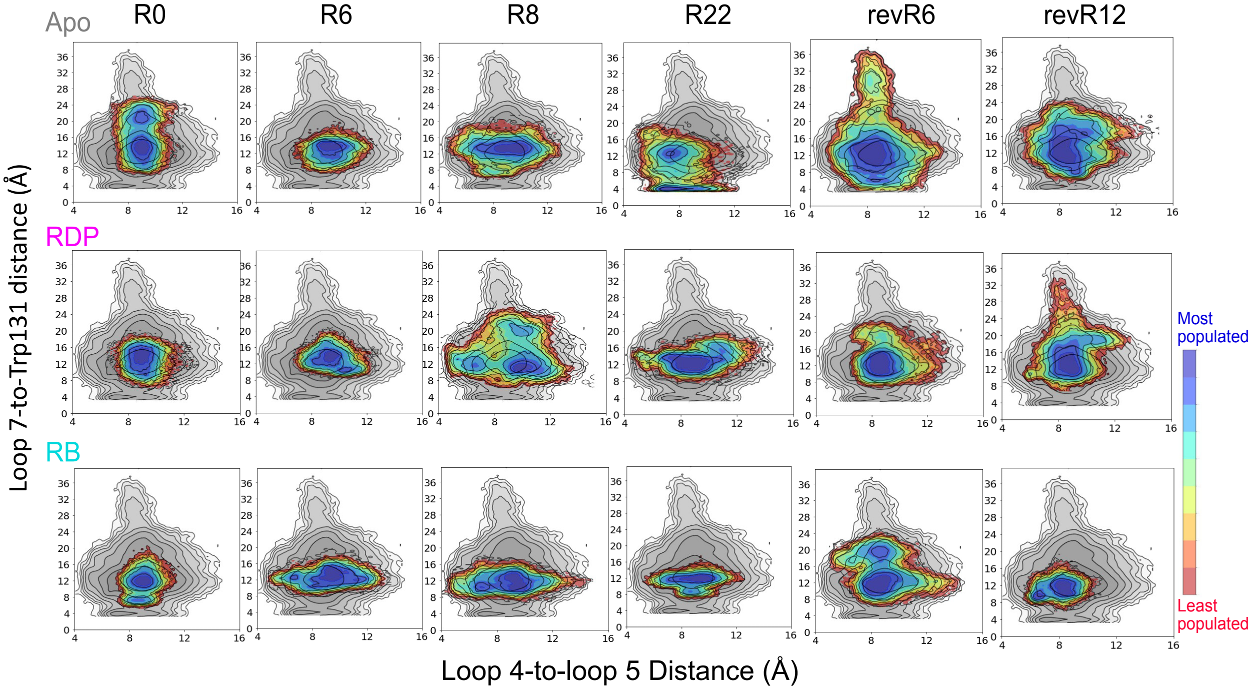


**Figure S15. 2D histograms of conformational dynamics in functionally important loop structures.** The histograms show loop 4-to-loop 5 distances and loop 7-to-Trp131 distances. Blue and red colors indicate the degree of the population in each state that the enzyme sampled. The gray color shows the entire conformational space that enzyme sample throughout evolution.

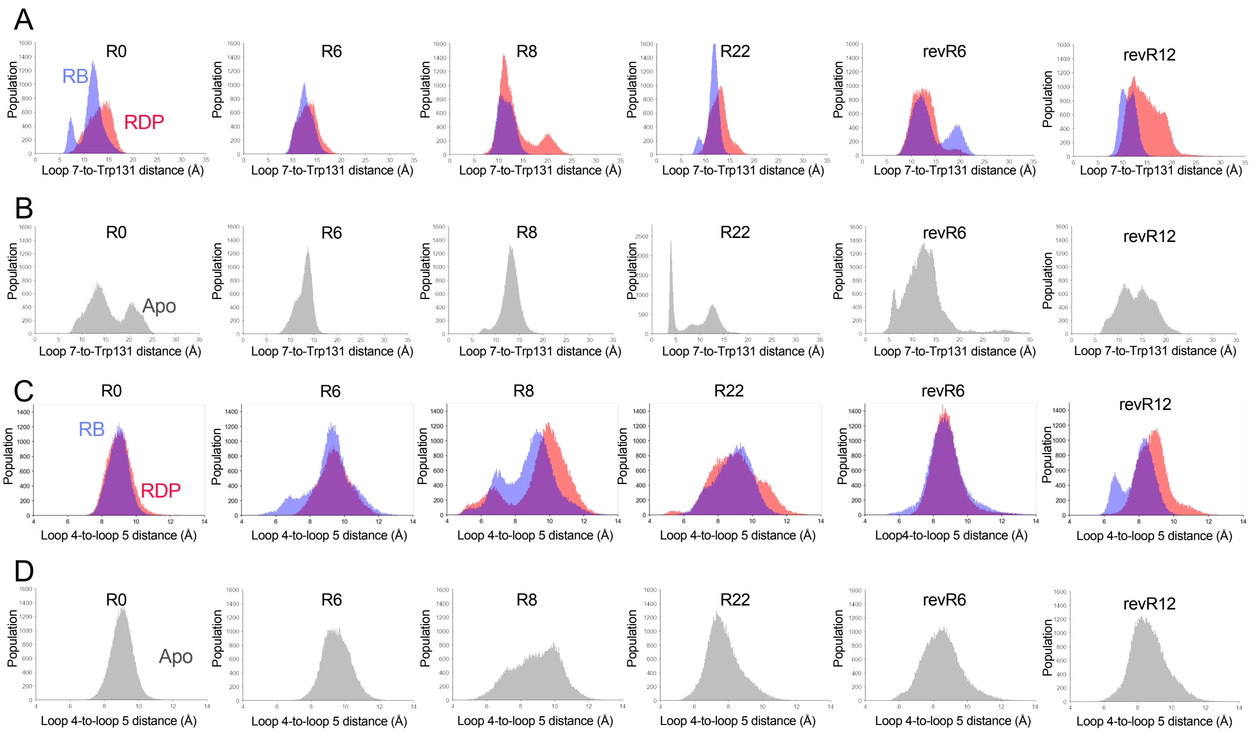


**Figure S16. 1D histograms of conformational dynamics in functionally important loop structures.** (A), (B) 1D histograms showing loop 7-to-Trp131 distances with substrates and apo-state. (C), (D) 1D histograms showing loop 4-to-loop 5 distances with substrates and apo-state.

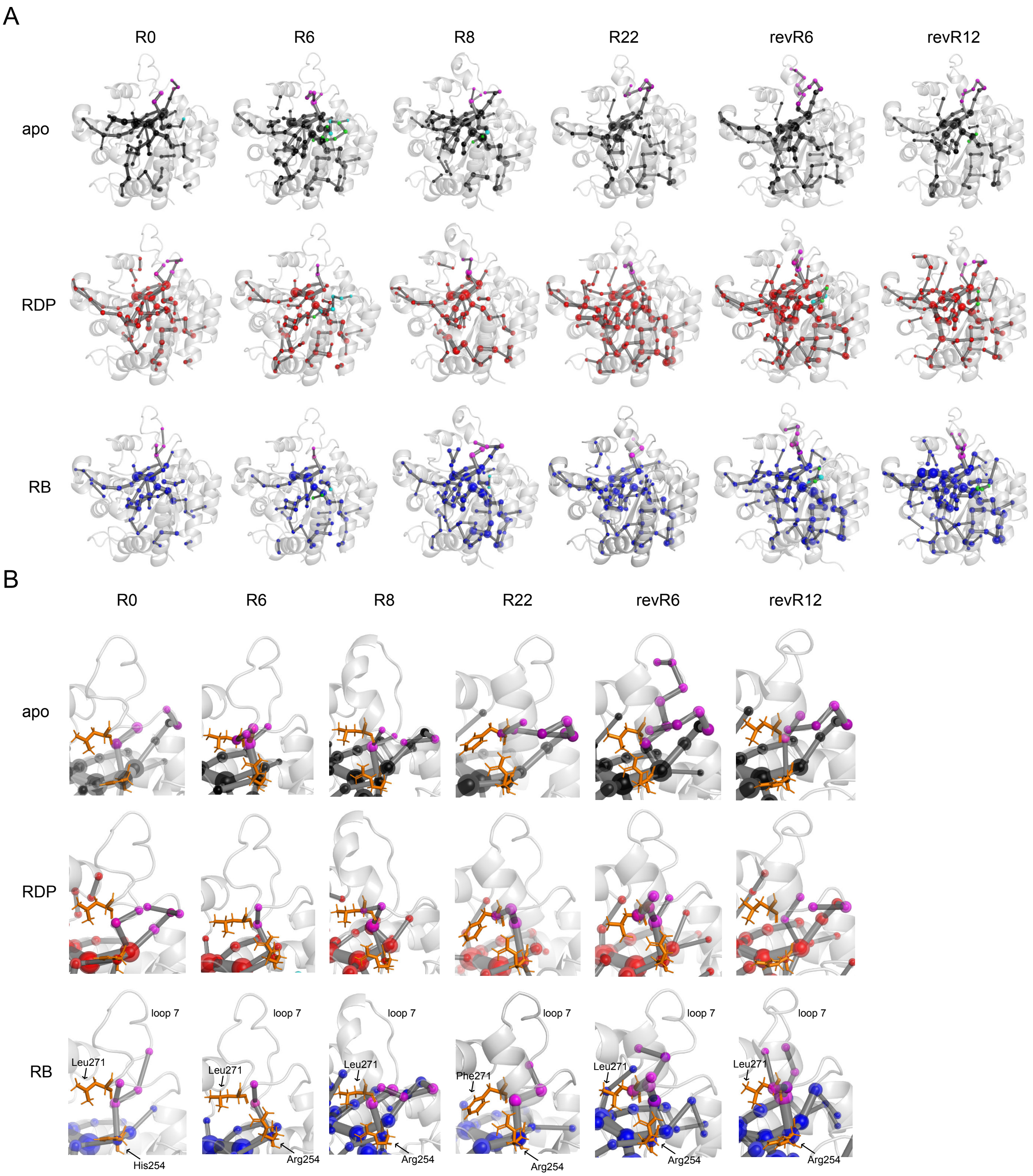


**Figure S17. Shortest path maps (SPMs) of PTEs.** (A) Alterations of SPMs in PTE-R0, 6, 8, 22, revR6, and revR12 were analyzed without substrate (apo state, black sphere) and with RDP (red sphere) and AE (blue sphere) substrates. Spheres and gray edges indicate correlation dynamics between the pair of residues. SPMs in loops 4, 5, and 7 are shown by spheres with green, cyan, and magenta. (B) Alteration of SPMs in loop 7.

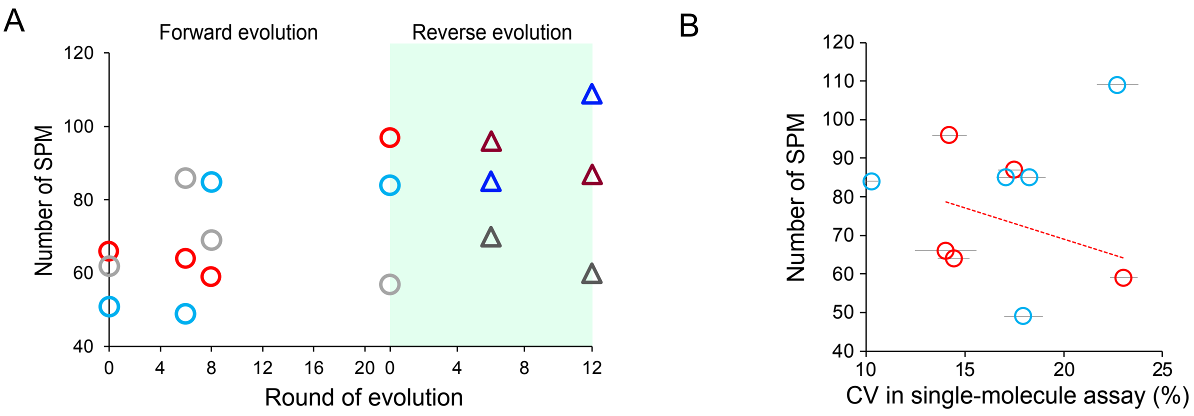


**Figure S18. The changes of SPMs in PTE and AE activities.** (A) The change in the number of SPMs measured with RDP (red circles and triangles) and RB (blue circles and triangles) substrates. Forward and reverse evolutions are shown as circles and triangles, respectively. (B) The correlation between *CV* in the single-molecule kinetic assay and the number of SPMs measured with RDP (Red circles) and RB (blue circles). The linear fittings of the data points in PTE activity provide fitting slopes; -161 (Pearson's correlation coefficient = 0.4)

### Supplementary Methods

#### Expression and purification of PTEs

Genes encoding PTEs were cloned into the pET-Strep vector used in previous studies [35, 36]. The pET-Strep-tag-PTE plasmid was transformed into BL21DE3 cells by electroporation. The transformed cells were pre-cultured overnight at 37°C in 5 mL of LB medium containing 100 µg/mL ampicillin. For protein expression, 0.5 mL of the bacterial culture was transferred to 10 mL of Overnight Express Instant TB medium (MilliporeSigma, Burlington, MA, USA) supplemented with 100 µg/mL ampicillin and 200 µM $ZnCl_2$ in a 50 mL tube. Bacteria were cultured at 30°C for 8 h, and then protein expression was induced at 16°C for 12 h. Cells were harvested by centrifugation and stored at -80°C for at least 1 day. We followed a previously developed method for lysate preparation [36]. Cells were lysed with 2 mL of a lysis buffer (1:1 mixture of B-PER and 50 mM Tris-HCl (pH 8.5) containing 100 mM NaCl, 200 µM $ZnCl_2$, 100 µg/mL lysozyme, and 0.2 µL of benzonase). After incubation for 1 h at room temperature, cell debris was spun down, and lysate containing expressed protein was collected. MagStrep beads (IBA Lifesciences, Goettingen, Germany) were used for purifying strep-tagged PTE. The beads were pre-equilibrated with wash buffer 1 (50 mM Tris-HCl (pH 8.5) containing 100 mM NaCl and 200 µM $ZnCl_2$) and mixed with the lysate for 30 min on ice with shaking. After binding, the beads were washed three times with wash buffer 1 and wash buffer 2 (50 mM Tris-HCl (pH 7.5) containing 100 mM NaCl, 200 µM $ZnCl_2$). The protein was eluted with 50 mM biotin dissolved in wash buffer 2 by incubating it on ice with shaking for 10 min. Finally, the eluted protein solution was desalted by using a spin desalting column (Bio-Gel, Bio-Rad, Hercules, CA, USA) equilibrated with wash buffer 2. The collected protein solution was stored at 4°C until use.

For large-scale expression and purification, 200 mL of Overnight Express Instant TB medium was prepared in a 2L flask. 30 mL lysis buffer per 8 g bacterial pellet was used for lysate preparation. The cell lysate passed over a Strep-Tactin column (IBA Lifesciences) with a 2 mL resin volume for protein binding. The column was pre-equilibrated with 6 mL of wash buffer 1. After binding, the column was washed with 6 mL wash buffer 1 and wash buffer 2. Then, the protein was eluted with 10 mL of 50 mM biotin solution. After elution, the protein solution was desalted using a desalting column (Econo-Pac 10DG, BioRad, Hercules, CA, USA), and the protein solutions were concentrated using a spin filter (Spin filters 10k, Pall Laboratory, Port Washington, NY, USA).

**Analyzing monomer-dimer transition in PTE molecules.**

We assessed the possible monomer-dimer transition of PTE molecules with fast protein liquid chromatography (FPLC) and Blue-native PAGE. We used FPLC (Duoflow, Bio-Rad) equipped with a size exclusion column (Superdex 200 5/150GL, Cytiva, Marlborough, MA, USA). 1.0 mg/mL protein solution was loaded onto the FPLC, and the flow rate was set to 0.1 mL/min. A gel filtration standard (Biorad) with FPLC prepared a calibration curve for estimating molecular weight.

Coomassie brilliant blue G-250 (Thermo Fisher Scientific Inc.) was used for Blue-native PAGE to stabilize complexes [44]. The protein solution was mixed with a sample loading buffer (62.5 mM Tris-HCl (pH7.0), 40% w/w glycerol, 0.01% w/w bromophenol blue, 0.02% w/w Coomassie brilliant blue G-250) in a 1:1 ratio. The sample was loaded onto a gel (4-20% Mini-protein TGX precast gel, Bio-Rad) and subjected to gel electrophoresis using a running buffer (25 mM Tris, 192 mM Glycine, 0.01% w/w

Coomassie brilliant blue G-250). Gel was immersed in a destaining buffer (30% v/v Methanol, 10% v/v Acetic acid) for one day. We run both PTEs and purified dimeric *E.coli* alkaline phosphatase (86 kDa) as positive controls with 1.0 and 3.0 mg/mL.

**Fabrication of microreactor array device**

The microreactor array device was fabricated with photolithography and etching [32, 34, 39]. The cover glass (24-40 mm, thickness 0.13-0.17 mm, Millipore Sigma) was incubated in 10 M KOH solution (Thermo Fisher Scientific Inc., Waltham, MA, USA) for 30 min. After washing with deionized water and drying, hexamethyldisilazane was coated onto the glass at 3,000 rpm for 30 s using a spin coater. After 5 min baking at 98°C on a hot plate, fluoroacrylic polymer (FluoroPel, CYTONIX) was coated onto the glass at 3,000 rpm for 30 s. The glass was then baked at 98°C for 5 min. A positive photoresist (AZ-4903, Merck KGaA, Darmstadt, Germany) was coated onto the glass at 500 rpm for 10 s then 8,500 rpm for 40 s. The glass was baked at 98°C for 5 min. Photolithography was then performed using a mask aligner (NxQ4006 aligner, Neutronix-Quintel, Morgan Hill, CA, USA) with a chrome photomask containing a 4 μm holes array. The photoresist on the glass was then developed using a photoresist developer (AZ300 MIF, Merck KGaA) for 5 min. The patterned glass coverslip was washed with deionized water. After drying, the glass was etched with $O_2$ plasma using a reactive ion etching machine (PE-50, Plasma-Etch, Carson City, NV, USA). Then, the remaining photoresist on the glass was entirely removed using 2-propanol and ethanol, and the glass was rinsed with deionized water. The diameter and thickness of the reactor were measured using a 3D interferometer

(Polytec, Irvine, CA, USA). The diameter and thickness of the reactor array used in these experiments were approximately 5.0 μm and 100 nm, respectively.

**Chemical synthesis of resorufin-diethylphosphate (RDP)**

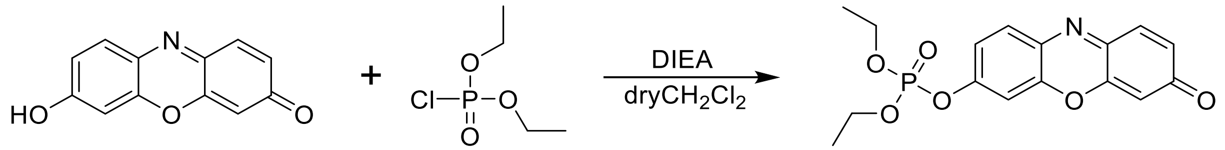


Resorufin (200 mg, 0.9 mmol, Biosynth, Staad, Switzerland) and diethyl chlorophosphate (174 mg, 1.1 mmol, Tokyo Chemical Industry Co., Ltd., Tokyo, Japan) were dissolved in dry dichloromethane (15 mL) with DIEA (235 μL, 1.3 mmol, Tokyo Chemical Industry Co., Ltd.). The reaction mixture was stirred for 6 h at room temperature. Reaction progress was followed by TLC (silica gel 60 $F_{254}$, Merck, $CHCl_3$: EtOAc = 3: 2). The reaction mixture was washed with 0.1M HCl (50 mL, 2 times) and saturated NaCl (50 mL, 1 time). The organic phase was dried with $Na_2SO_4$, filtered, and the solvent was evaporated. The product was purified by silica gel column chromatography with a gradient of 0% to 40% EtOAc in $CHCl_3$ to yield the target product (60 mg, 19% yield) as an orange solid. $^1$H NMR spectra were recorded on a JNM-ECP300 FT NMR spectrometer (JEOL Ltd., Tokyo, Japan) and chemical shifts were expressed in parts per million (ppm), and the coupling constants were calculated in Hz ($^1$H NMR (300 MHz; $CDCl_3$): $\delta$ 7.78 (*d*, 1H, $J_H$ = 9.3 Hz, Ar-H), 7.42 (*d*, 1H, $J_H$ = 9.9 Hz, Ar-H), 7.26 (*m*, 2H, Ar-H), 6.85 (*d*, 1H, $J_H$ = 9.9 Hz, Ar-H), 6.33 (*s*, 1H, Ar-H), 4.27 (*m*, 4H, -$CH_2$-Me), 1.39 (*t*, 6H, $J_H$ = 7.1 Hz, -$CH_3$)). ESI-TOF-MS spectra were recorded on a JMST100LP spectrometer (JEOL) (ESI-TOF-MS *m/z* = 350.09 $[M+H]^+$ (calcd. for $C_{16}H_{17}NO_6P$, 350.08)).

**Single-molecule assay**

Reaction buffer (100 mM Tris-HCl (pH8) supplemented with 200 μM $ZnCl_2$, 0.1% Triton X-100, 25 μg/mL bovine serum albumin (MilliporeSigma), 20 μM Resorufin) was used for single-molecule kinetic assay. Resorufin-diethylphosphate (RDP), resorufin-butyrate (RB, MilliporeSigma), and resorufin dye (MilliporeSigma) were dissolved in dimethyl sulfoxide (DMSO) and stored at -20°C until use.

The flow channel was constructed on the surface of the microreactor array device using double-sided adhesive tape (CAPLINQ, Assendelft, Netherlands) and a top cover made of polydimethylsiloxane (PDMS, Dow Corporate, Midland, MI, USA). The silicone elastomer and curing agent were mixed at a 1:10 ratio using a mixer (Thinky, Tokyo, JAPAN). After being degassed for 30 min, PDMS was solidified on a petri dish at 60°C for 1 day. PDMS was cut to the size of the device and the inlet and outlet holes were punched with a puncher. The double-sided tape was cut out in the shape of the flow path and was applied to the surface of the device. Then, the PDMS lid was attached on top of the double-sided tape.

For the single-molecule kinetic assay, the diluted enzyme (final ~5 pM) and reaction buffer were first mixed, followed by the addition of the fluorogenic substrate (final 200 μM), just before applying the reaction mix to the flow channel. After introducing the reaction mix, air was applied from the same inlet to remove the excess solution and seal the reactor. Measurements began 30 seconds after sealing. A new flow channel was used for each assay. The same setup as described above was used in the reaction solution exchange experiment. To prevent x-y drift during sample loading, the device was fixed on the microscope stage, and the reaction solution was applied using a gel-loading chip (VWR, Radnor, PA, USA). After measuring the first reaction, the device was washed twice with 200 μL of fresh reaction buffer, except for the substrate. Following a 10-minute incubation to ensure the complete removal of the reactants, the second reaction was measured by applying the reaction buffer with the substrate.

All images of single-molecule assays were acquired using an inverted fluorescence microscope (Eclipse Ti-E, Nikon, Tokyo, Japan) equipped with a 40× objective lens (Nikon, Plan Apo, NA = 0.9), multi-wavelength LED illumination (SpectraX, Lumencor, Beaverton, OR, USA), EM-CCD camera (imagEM, C9100-13, Hamamatsu Photonics, Shizuoka, Japan), an incubator for temperature control, and a Perfect Focus System (Nikon). The fluorescence of the fluorescein dye was measured using filter sets (excitation 550/15 nm, dichroic 573 nm, emission 598/25 nm, Nikon). All measurements were performed using the NIS-Element software (Nikon). The temperature of the incubator was set to 22°C. Time-course images were taken at 2 min intervals for 10 min. The cross-reactor diffusion of the fluorescein dye was evaluated by photobleaching using a field diaphragm (Nikon) to limit the area of excitation light (**Fig. S3**). All images of single-molecule assays were analyzed using a custom Python code [34].

**Analysis of single-molecule data.**

The single-molecule activity was determined by plotting the histogram of the fluorescence intensity distribution at the end-point (**Fig. 2C**). We typically observed an enzyme-free population (peak 0), a bimodal distribution next to peak 0 (peak 1L and peak 1H), and an additional minor peak (peak 2). Firstly, we fitted the peak 0 with a single Gaussian function, and the mean intensity ($MI_0$) and standard deviation ($SD_0$) were calculated. Enzyme activity was observed above a threshold value defined as $MI_0 + 10 \times SD_0$. Then, we statistically identified peak 2 and removed it from the data to accurately analyze single-molecule activity [32, 34, 39]. After removing peak 0 and peak 2, the time trajectories of enzyme molecules were linearly fitted, and slopes (a.u./min) were calculated. Slopes can be converted to the turnover rate ($s^{-1}$) by using a calibration curve that provides the concentration of the resorufin as a function of fluorescence intensity (**Fig. S5B**). After the conversion and subtracting autohydrolysis rates of RDP and RB substrates (16 and 26 ($s^{-1}$)), the distribution of the turnover rate was plotted (**Fig. S4**).

The activity distribution, or functional substates, of each peak, was calculated as $CV_{1H \text{ or } 1L}$ (%) = $SD_{1H \text{ or } 1L}/MT_{1H \text{ or } 1L} \times 100$. For calculating the overall functional substates of peak 1L and peak 1H, the sum of $CV$ ($CV_n$) was calculated with the following equation;

$$CV_n = \sqrt{CV_{1L}^2 + CV_{1H}^2}$$

The number of analyzed enzyme molecules was higher than 700 in all measurements (see **Table S1**).

**Analysis of the reaction solution exchange experiment**

To calculate the correlation among different functional substates in the reaction solution exchange experiment, we first extracted single-molecule activities from both two independent reactions in the density scatter plots (**Fig. S8A**). We defined threshold values using mean turnover rate ($MT_{1st/2nd\_1L \text{ or } 1H}$) and standard deviation ($SD_{1st/2nd\_1L \text{ or } 1H}$) of peak 1L and peak 1H observed in the 1$^{st}$ and 2$^{nd}$ reactions. The threshold values for the two reactions were calculated as $MT_{1st/2nd\_1H} + 2 \times SD_{1st/2nd\_1H}$ and $MT_{1st/2nd\_1L} - 2 \times$

$SD_{1st/2nd\_1L}$, respectively (red broken lines in **Fig. S8A**). Then, the population inside the red broken lines was further classified into four clusters. For classification, the local minima between peak 1L and peak 1H were calculated in both reactions (cyan broken lines in **Fig. S8A**), and the four clusters were classified based on the cyan broken lines through the points. Enzyme molecules belonging to clusters 1 and 2 were those that belonged to the same active peak in the two measurements, while molecules belonging to clusters 3 and 4 were those that transitioned between the active peaks.

The repeatability of the reaction solution exchange experiment was assessed by using resorufin dye (**Fig. S8B**). Resorufin dye was encapsulated in the reactor, and the fluorescence intensity of around 10,000 reactors was taken. Then, the dye was washed out and fresh dye was encapsulated in the same reactors. We repeated the encapsulation/wash process four times and calculated the mean fluorescence intensity and *CV* of the distribution of the fluorescence intensity in the reactors. The upper detection limit of the reaction solution exchange experiment was also analyzed by using the resorufin dye. We measured the distribution of resorufin fluorescence intensity twice by performing the reaction solution exchange and calculated Spearman's correlation coefficient between 1$^{st}$ and 2$^{nd}$ reactions.

**Assessment of double occupancy in the activity distribution.**

While nearly all PTEs exhibited bimodal distributions (peak 1L and peak 1H), there is a possibility that peak 1H arises from the double occupancy of enzyme molecules associated with peak 1L. Therefore, we statistically assessed the single and double occupancy of enzyme molecules in the reactor. As discussed in our previous study [32,

34, 39], the number of enzyme molecules in a reactor follows a Poisson distribution. In a Poisson distribution, the probability of a given number of enzyme molecules can be calculated as follows:

$$P = \frac{e^{-\lambda}\lambda^{k}}{k!}$$

where $k$ is the actual number of enzyme molecules in a reactor, and $\lambda$ is the expected mean number of enzyme molecules per reactor. $\lambda$ can be calculated by dividing the number of reactors exhibiting enzyme activity by the total number of observed reactors.

In the case of PTE-R22 (**Fig. 1C**), $\lambda$ was determined to be 0.56, and we expected approximately 32% of the reactors to contain single enzyme molecules, 11% to contain multiple enzyme molecules (mainly 2-3), and 57% of reactors to be enzyme-free. As a result, the ratio of single occupancy to multiple occupancy of enzyme molecules was around 3. The experimental ratios of single, multiple, and enzyme-free were 32%, 7%, and 61%, respectively. Thus, the ratio of single occupancy to multiple occupancy was around 5, which agrees with Poisson statistics. Assuming that peak 1H results from double occupancy of peak 1L, the experimental population ratio of peak 1L to peak 1H is 0.83, which is significantly different from the expected ratio of 3, indicating that peak 1H was not caused by double occupancy of peak 1L.

PTE-R2a exhibited a higher population in peak 1L than in peak 1H (**Fig. S4**). In this case, experimental $\lambda$ was determined to be 0.26, and we expected approximately 20% of the reactors to contain single enzyme molecules, 3% to contain multiple enzyme molecules, and 77% of the reactors to be enzyme-free. Therefore, the ratio of single occupancy to multiple occupancy was around 7. The experimental percentages of single, multiple, and enzyme-free were 76%, 21%, and 3%, respectively, and the ratio was around 7, which agrees with Poisson statistics. However, the population ratio of peak 1L to peak 1H is 1.4, which is significantly lower than the expected ratio of 7 if peak 1H were due to the double occupancy of peak 1L. This suggests that peak 1H is not a result of double occupancy. Based on these results, the bimodal distribution observed in peak 1 was not due to the multiple occupancy of the enzyme molecule but rather the enzyme having two distinct activity states. We identified peak 2 in the same manner, and since peak 2 appears at approximately twice the mean turnover rate of peak 1, we omitted peak 2 from the analysis of peak 1 near the local minima of the overlap between peak 1 and peak 2 (black broken lines in **Fig. 2C**). Otherwise, we defined peak 1 as a bimodal distribution.

**Assessment of the accuracy of single-molecule assays**

The intrinsic noise derived from the microscope system and variation in reactor volume can be estimated from the distribution of fluorescence intensity in reactors encapsulating the resorufin dye. The distribution of the fluorescence intensity across over 20,000 reactors was analyzed, and the coefficient of variation ($CV_{\mathrm{reso}} = 100 \times SD_{\mathrm{reso}}/MI_{\mathrm{reso}}$) was calculated. The $CV_{\mathrm{reso}}$ ranged from 0.3 to 3.3% at resorufin dye concentration from 10 to 400 μM (**Fig. 5B**). The $CV$ of peak 0 ($CV_0$) in all measurements was 0.7 ± 0.4% (mean ± $SD$), indicating that all measurements were performed with high accuracy, very close to the detection limit of heterogeneity. The noise derived from time-course measurements was assessed by measuring the autohydrolysis of the substrates.

We evaluated the potential dynamic transition of catalytic activities. The coefficient of variation of fluorescence intensity, calculated from two successive time points

($CV_{fluctuation}$ = $SD$ ($I_t$, $I_{t-1}$)/mean ($I_t$, $I_{t-1}$), where $SD$ and mean are calculated from the fluorescence intensity at time t), was calculated with time-course trajectories of PTE-R22 (**Fig. 1C**). The $CV_{fluctuation}$ of R22 was 2.0 ± 0.5 (%), which is low enough to be negligible, indicating that observed activity state of PTE is stable within the observation period, similar to other enzymes such as β-galactosidase, β-glucuronidase, and alkaline phosphatase, reported previously [30-34, 39]. The reaction solution exchange experiment also supports a less dynamic transition (**Fig. 3B-C** and **Fig. S9**).

The effect of enzyme molecule adsorption onto the reactor surface was verified by performing reaction solution exchange experiments (**Fig. 3B-C** and **Fig. S9**). We identified adsorbed and escaped enzyme molecules at 2$^{nd}$ reaction and analyzed the turnover rate and $CV_n$ of those populations. Positive linear correlations of the turnover rates between adsorbed and escaped molecules suggest that the binding of enzyme molecules on the reactor surface did not change their kinetic functions (**Fig. S6B**).

**Bulk ensemble kinetic assay**

Bulk ensemble kinetic assay of PTE was performed in a 384-well black plate using a plate reader (Synergy H1, BioTek, Winooski, VT, USA). For measurements with the resorufin dye, the excitation and emission wavelengths were set to 550 nm and 590 nm, respectively. Changes in fluorescence intensity were acquired at thirty-second intervals at 26°C. A reaction solution (50 µL) was prepared by mixing PTE and substrate in the reaction buffer (100 mM Tris-HCl (pH 8) supplemented with 200 µM $ZnCl_2$, 0.1% Triton X-100, 25 µg/mL bovine serum albumin (MilliporeSigma)), and the mixture was then applied to the plate wells. Three independent measurements were taken to analyze the mean and

*SE* of enzyme activity. Enzyme activity was determined from a linear fit of the increase in fluorescence intensity over time and converted to the turnover rate with a calibration curve (**Fig. S5A**). The initial rate was measured using six different concentrations and fitted to the Michaelis-Menten equation.

**Calculation of β-factor from crystal structures**

We calculated the β-factor using the previously solved crystal structures of R0 (4PCP), R2 (4XD5), R6 (4XAG), R8 (4XAY), R18 (4XAZ), R22 (4PCN), revR6 (4PBE), and revR12 (4PBF). The β-factor of the Cα atoms was normalized by the mean Cα β-factor of the entire chain in each subunit. We then collected the normalized β-factors for loop 4 (residues 170-175), loop 5 (residues 200-210), and loop 7 (residues 257-275) and calculated the average β-factor across subunits.

**MD simulations of PTEs**

Several PTEs, including PTE-R0, R6, R8, R22, revR6, and revR12, were simulated using the Amber18 package, with both substrate-free and substrate-bound states (RDP and RB) [45]. Structures of PTE were obtained from the RCSB Protein Data Bank with the following PDB IDs: 4PCP (R0), 4XAG (R6), 4PBE (revR6), 4XAY (R8), 4PBF (revR12) and 4PCN (R22). The structure of resorufin butyrate (RB) was obtained from the PubChem database (CID 500716). Diethylphosphate group was added to the resorufin (CID 69462) to create resorufin diethylphosphate (RDP) using ChemDraw and further optimized with Gaussian09 (see below). These two substrates were used to superpose onto the naphthalene ring of the transition state analog of the AE substrate bound to R18

(PDB ID: 4E3T) to determine the binding positions of RDP and RB. Further, these substrates were superimposed onto both chains A and G. Topology and parameters of the substrates were generated using the Gaussian09 software package by utilizing B3LYP/6-31G* level of theory. The protonation states of these complexes were corrected using H++ server [46]. Further, these systems were prepared using the tleap program to generate a system with the protein inserted in a cubic water box of 15 Å. The system was neutralized using counter ions. One OH group was position-restrained with the two Zn atoms at a distance of 2 Å. Additionally, the OH group of ligands was distance-constrained with one of the Zn atoms. The simulation was performed for the apo and holo (RB and RDP) structures. Minimization was performed for 10,000 steps. The heating process was performed stepwise by increasing the temperature from 50K to 300 K. Equilibration and production runs were performed at 300 K. The equilibration process was done in two steps for 20,000 steps. These equilibrated structures were simulated using Gaussian-accelerated molecular dynamics for 500 ns which adds a harmonic boost potential to enhance conformational sampling. A dual boost for dihedral energy and potential energy was applied, with the threshold energy set to the upper bound. The upper limit of the standard deviation for the dihedral and total energy boost potentials was set to 6.0 kcal/mol. First, a 10 ns conventional MD simulation was run to calculate the GaMD acceleration parameters. In a second step, a 50 ns GaMD equilibration run was performed. The average and standard deviation of boost potentials were collected every 0.5 ns. Furthermore, a 500 ns GaMD production run was performed in three independent replicas. Similarly, the apo structures were also simulated for 500 ns (in three replicas) using the Amber18 program, with the ff14SB force field and TIP3P water model [46] used as the solvent.

Shortest path maps (SPMs) were calculated considering for the Cα atoms of the protein residues along the trajectory. The distance and correlation matrix were used for building the SPM networks, which displays the pairs of nearby (within 6 Å) residues that are highly correlated during the molecular dynamics trajectory [20, 21, 42, 43]. The total number of Structures were analyzed using MDTraj and PyTraj and visualized in PyMOL. The total number of edges represents the combined edges observed in the two subunits.

Conformational landscapes were calculated based on loops 4, 5, and 7 distances. The median (*Med*) and median absolute deviation (*MAD*) were calculated, and the coefficient of variation (*CV*, %) was determined by dividing *MAD* with *Med*.
